# Ancient persistence and recurrent emergence of structural variants across divergent Atlantic salmon lineages

**DOI:** 10.64898/2026.05.25.727466

**Authors:** Célian Diblasi, Jun Soung Kwak, Domniki Manousi, Mariann Arnyasi, Arturo Vera Ponce De Leon, Nicola Jane Barson, Marie Saitou

## Abstract

Structural variants (SVs) are a major source of genomic diversity, yet the evolutionary origins of SVs shared across divergent populations remain difficult to resolve. Shared SVs may reflect ancient polymorphism, recurrent mutation, introgression, or subsequent lineage-specific frequency change, but the relative contribution of these processes often remains difficult to distinguish.

Here, we investigated SV evolution across four Atlantic salmon (*Salmo salar*) lineages differing in geography, Europe versus North America, and domestication status, wild versus farmed. Using sensitive SV discovery, stringent genotyping, local PCA, haplotype-distance analyses, and forward simulations, we tested whether broadly shared SVs behave as a single class of variation or separate into distinct evolutionary categories. We generated a high-confidence SV map and found that SVs were enriched in repetitive regions, particularly segmental duplications and LTR retrotransposons, consistent with genome architecture shaping SV formation.

Nearly half of high-confidence SVs were shared across all four lineages despite deep continental divergence, and simulations showed that this broad sharing is more consistent with ancient persistence than recurrent mutation alone. In contrast, a small subset of large SVs exhibited complex PCA clustering and multimodal haplotype-distance distributions, consistent with recurrent formation at structurally unstable loci. Large SVs also showed contrasting frequency trajectories between continents, and one immune gene-rich copy-number variable region showed a marked frequency increase in domesticated European salmon. Together, these results show that shared SVs comprise distinct evolutionary categories shaped by ancient persistence, recurrent mutation, and lineage-specific frequency change.

## Introduction

Structural variants (SVs), including insertions, deletions, inversions, duplications, and translocations, are major sources of genomic diversity. Unlike single nucleotide polymorphisms (SNPs), which involve changes at individual base pairs, SVs can affect kilobase- to megabase-sized regions, altering gene dosage, regulatory landscapes, recombination, and chromosome structure. As such, SVs can have large effects on gene function, expression, and inheritance, and have been implicated in adaptation, reproductive isolation, and population divergence (Hämälä et al. 2021; Scott et al. 2021; Stuart et al. 2025)(Lowry and Willis 2010; Faria et al. 2019; Berdan et al. 2024; Ferguson et al. 2024; Hämälä et al. 2024).

A central unresolved question is how to interpret SVs that are shared across divergent populations. Such sharing may reflect ancient polymorphism, recurrent mutation at the same locus, introgression, or subsequent shifts in allele frequency caused by selection or demography. Distinguishing among these alternatives is difficult because SVs are often large, repetitive, and complex, and therefore challenging to genotype reliably with short-read sequencing. Although several studies have documented shared SVs across populations, fewer have distinguished whether such sharing reflects common ancestry, recurrent formation, or later redistribution through introgression or selection. (Lindtke et al. 2017; Wellenreuther et al. 2019; Aqil et al. 2023; Ferguson et al. 2024). Further, when structural variants are shared across populations, it remains unclear whether this pattern reflects recurrent mutation at the same locus, persistence of ancestral polymorphism across population splits, or selection shaping the frequency and distribution of these variants under either scenario.

Although short-read sequencing poses inherent challenges for SV discovery, recent advances in bioinformatic methods have improved the ability to detect and genotype SVs with higher confidence (The Danish Pan-Genome Consortium et al. 2018; Kosugi 2019). By applying such approaches, it is now possible to address long-standing questions about the origins and evolutionary dynamics of SVs even in species with large and complex genomes (Mérot et al. 2020; Berdan et al. 2021; Villoutreix et al. 2021; Yang et al. 2024).

When shared SVs are detected across populations, several non-mutually exclusive mechanisms may underlie them. These include the retention of ancestral polymorphism, particularly when divergence is recent, or long-term maintenance by balancing selection, which may be especially relevant for SVs due to potential heterozygote advantage (Pearse et al. 2019; Berdan et al. 2021). However, recurrent mutations may also arise in structurally unstable genomic regions, such as those enriched in transposable elements and segmental duplications (Boettger et al. 2016). These regions are known hotspots for SV formation: repeated sequences promote non-allelic homologous recombination (NAHR), while transposable element activity can introduce double-strand breaks and erroneous repair (Kidd et al. 2010; Carvalho and Lupski 2016). Hybridization or introgression can also transfer SVs across taxa (Almarri et al. 2020; Quan et al. 2021; Zhang et al. 2023). These mechanisms can generate similar presence–absence patterns across populations but predict different haplotype structures and frequency trajectories. Disentangling the relative contributions of these processes is critical for understanding how SVs shape evolutionary trajectories, yet few natural systems provide the necessary conditions for rigorous testing.

Atlantic salmon (*Salmo salar*) provides a well-suited system for addressing these questions. The species comprises two deeply diverged lineages, European and North American, that split roughly one million years ago and subsequently evolved largely in isolation, with evidence of postglacial secondary contact in some populations (Payne et al. 1971; King et al. 2007; Rougemont and Bernatchez 2018). Both lineages have also undergone independent domestication in recent decades, offering replicated contrasts between wild and aquaculture populations exposed to distinct selective pressures. In addition, major karyotypic variation, including chromosome fusions in the North American lineage, together with residual sequence homology from the salmonid-specific whole genome duplication, contributes to a structurally dynamic genomic background that may facilitate recurrent SV formation (Lubieniecki et al. 2010; Lien et al. 2016; Stenløkk et al. 2022).

In this study, we use Atlantic salmon to test whether SVs shared across divergent lineages behave as a single class of variation or instead comprise distinct evolutionary categories shaped by ancient persistence, recurrent mutation, and lineage-specific frequency change. We combine high-confidence SV genotyping, local haplotype-structure analyses, functional annotation, forward-time simulations, and haplotype-distance comparisons to evaluate alternative evolutionary scenarios. This framework allows us to move beyond describing SV sharing and to infer the processes by which shared structural variation originates, persists, and becomes differentiated among lineages.

## Results

### Construction of a high-confidence SV set for testing evolutionary origins of shared variation

To construct a high-confidence structural variant (SV) map, we analyzed four Atlantic salmon lineages comprising 368 individuals. These lineages differed by their geographical origins (American vs European) and their domestication status (Wild or domesticated). We first applied Manta, a graph-based SV caller that integrates paired-end and split-read signals to achieve high sensitivity, identifying a total of 302,843 putative SVs. While highly sensitive, Manta’s detection strategy may also capture variants with limited genotyping resolution. To improve accuracy in genotyping, which is particularly important for comparative analyses across lineages, we subsequently used BayesTyper, a k-mer-based probabilistic genotyper that jointly considers read support and variant presence probabilities.

This two-step approach, combining sensitive discovery with stringent probabilistic refinement, yielded a final set of 6,683 high-confidence SVs (hereafter, BayesTyper_SVs; **Supplementary Figure 1**), in line with previous short-read based study (Bertolotti et al. 2020). The substantial reduction in variant number reflects the conservative nature of BayesTyper’s genotyping criteria, which are designed to minimize false positives, particularly in repetitive or low-coverage regions. Importantly, comparative analyses of retained and discarded SVs revealed that key features such as size and type were broadly preserved (**Supplementary Figure 2**), indicating that the refinement process enhanced genotyping reliability without introducing strong biases.

To better understand the characteristics of the high-confidence SV set, we next examined their size distribution, genomic context, and population-level patterns. The size distribution of BayesTyper_SVs exhibited two distinct peaks, with 27% of variants falling within the 50-100 bp range (**Figure 1A**). A secondary enrichment was observed around 1,432-1,436 bp, consistent with previously reported signatures of the *Tc1* mariner transposable element (Bertolotti et al. 2020). Among the 92 SVs in this range, 83 showed >95% identity to the *Tc1* mariner reference (EF685967.1), corroborating earlier findings and supporting the technical robustness of our detection pipeline. In terms of genomic distribution, SVs were significantly enriched in intronic regions compared to exons or coding sequences (**Figure 1B**). This pattern aligns with the expectation that purifying selection constrains the maintenance of structural variation in functionally important genomic regions, a trend observed in other vertebrate genomes as well (Zhou et al. 2019; NHGRI Centers for Common Disease Genomics et al. 2020; Hämälä et al. 2021).

**Figure 1:**
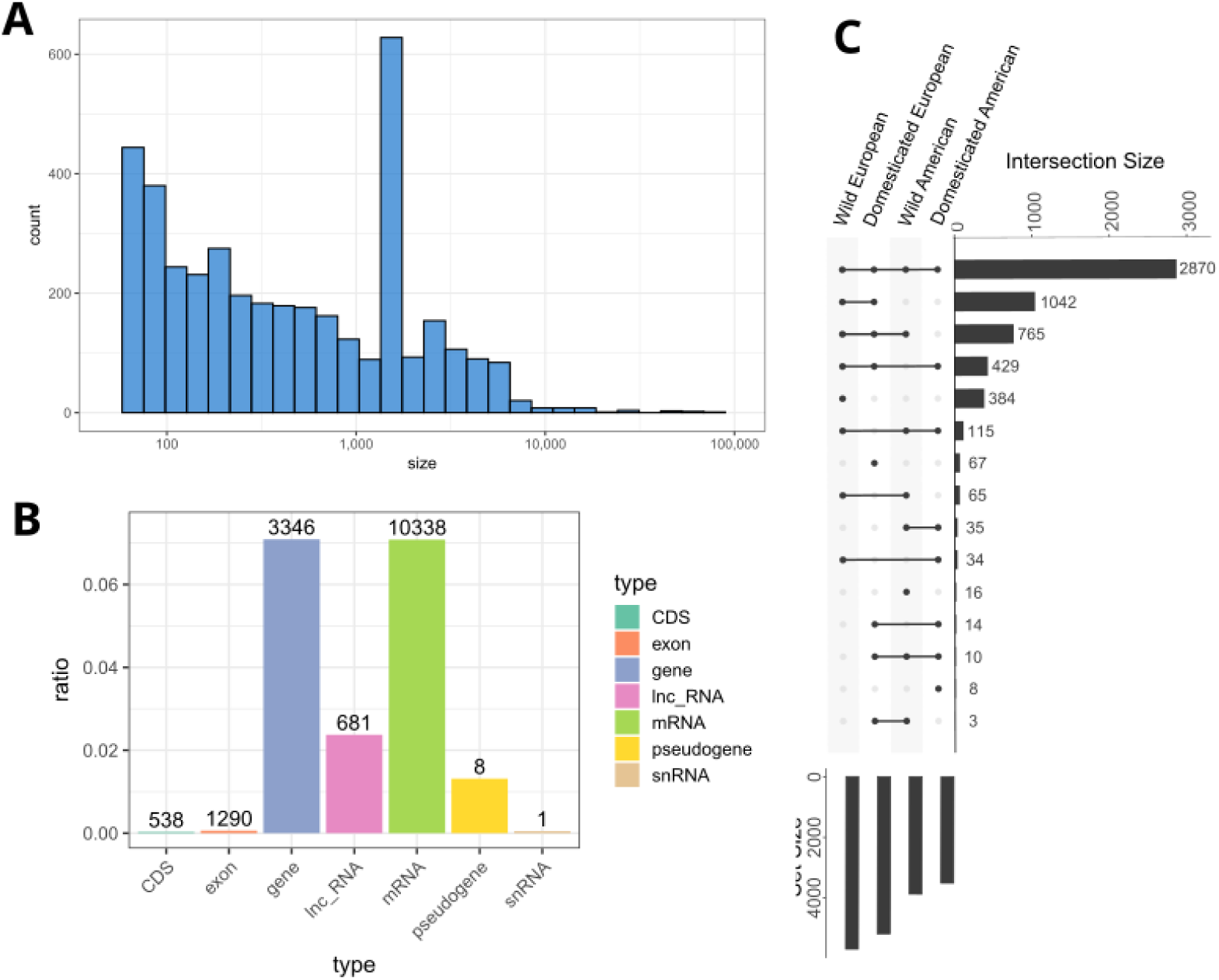
Overview of BayesTyper_SVs features. **A:** Size frequency plot of SVs in bp on a log10 scale. **B:** Proportion of SVs in genomic features, calculated for each annotation by taking the number of annotations overlapped for each type (indicated above the bars) divided by the total number of annotations. **C:** Upset plot of the SV sharing across the four lineages. Lineage names indicate their domestication status (Domesticated vs wild) and geographic origin (American vs European). Alt text: A: Bar plot of SV size showing higher proportion of small SVs as well as a peak around size 1500. B: Bar plot indicating the proportion of SVs overlapping different genomic features, with gene and mRNA the most overlapped annotations and CDS, exon and snRNA the least overlapped. C: Upset plot for SVs among the four lineages, showing that he most present category are SVs present in the four lineages

### SVs were enriched in repetitive regions, particularly LTR retrotransposons

To explore genomic features that may facilitate the formation of SVs, we assessed the extent to which BayesTyper_SVs overlapped with repetitive elements. Specifically, we focused on the overlap of SVs with segmental duplications and transposable elements, both known to promote genome instability through mechanisms such as non-allelic homologous recombination (NAHR). Our analysis revealed that SVs overlapped with repetitive sequences at a significantly higher rate than expected by chance (**Figure 2**). SVs overlapping segmental duplications were particularly enriched, ranking in the top 0.5% of a null distribution generated from random genomic locations. This enrichment is consistent with the well-known role of segmental duplications in facilitating NAHR. This observation suggest that segmental duplications often coincide with genomic areas where SVs arise more frequently (Escaramís et al. 2015; Carvalho and Lupski 2016).

**Figure 2:**
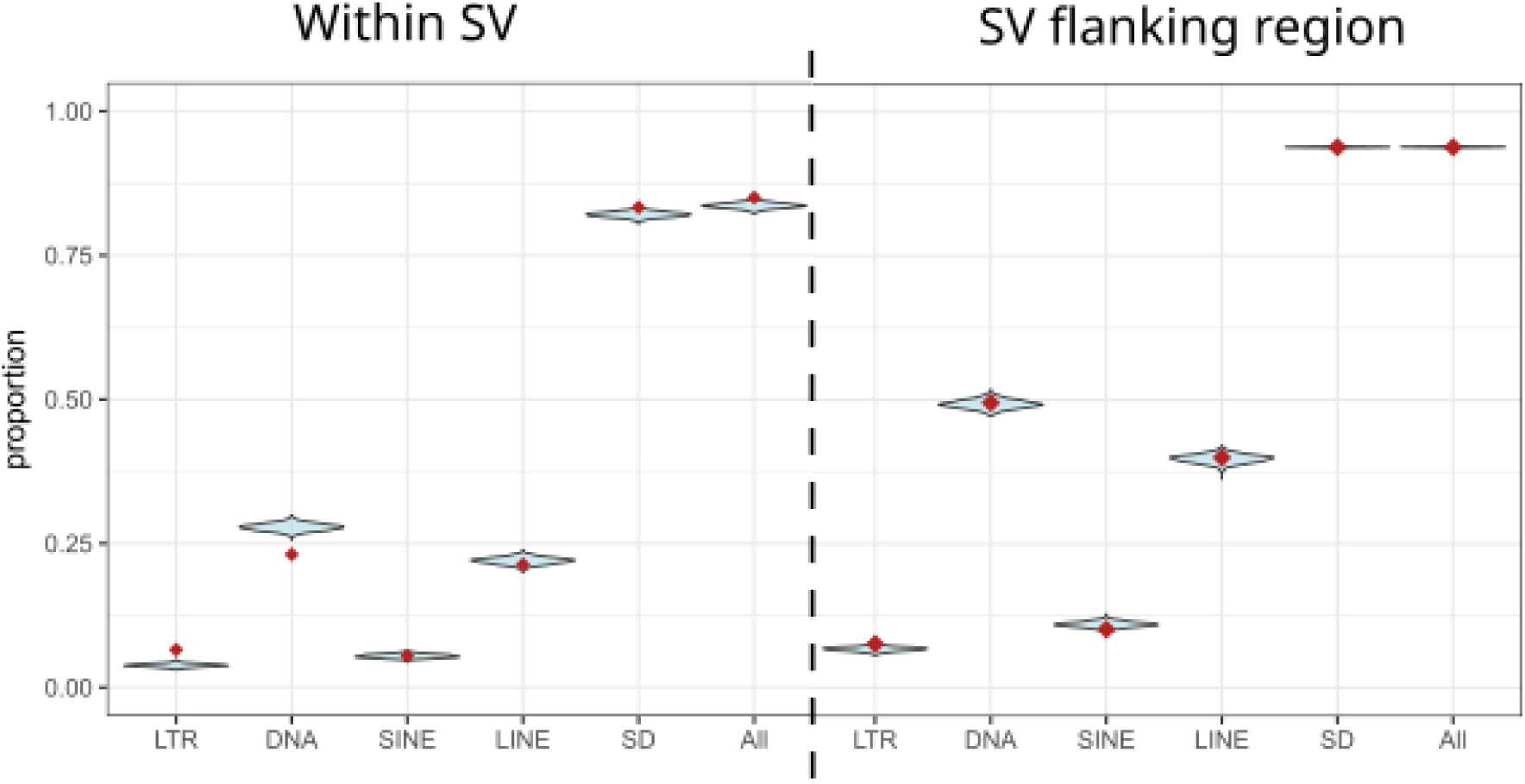
Proportion of SVs overlapped or surrounded by repeats compared to random expectations. The red dot indicates the observed proportion of BayesTyper_SVs overlapping (or surrounding) the given repeat type. The pale blue distribution shows the corresponding proportions obtained from 1,000 sets of randomly sampled genomic intervals matched to the observed SVs in number, size, and chromosomal distribution. Left: SVs overlapping repeats, right: SV surrounding repeat. Note that one SV can overlap or surround several repeat types. Alt text: Dot indicating the proportion of SVs overlapped by different annotation, together with violin plot of expected overlap in random genomic regions.

We further found that SVs were strongly enriched within or near LTR retrotransposons, both in terms of direct overlap and within 5 kb of flanking regions. In contrast, DNA transposons showed a significant depletion of overlapping SVs. These contrasting patterns likely reflect fundamental differences in transposition mechanisms: LTR retrotransposons propagate via a “copy-and-paste” mechanism and often introduce larger structural changes, while DNA transposons use a “cut-and-paste” strategy, which is less likely to promote extensive rearrangements (Gray 2000; Burssed et al. 2022).

### Nearly half of SVs were shared across all four lineages

Principal component analysis (PCA) based on genome-wide SVs largely recapitulated known population structure from SNP data, showing clear continental differentiation and a domestication-related separation within Europe (**Supplementary Figure 3A**; Buso et al.), suggesting that many SVs are neutral. One individual of our dataset, however, displayed a highly divergent haplotype structure and was consistently isolated in PCA space across multiple chromosomes. Based on this consistent outlier pattern, we excluded the individual from downstream analyses to avoid technical distortion in population-level analysis.

Analysis of SV sharing revealed that 49% of BayesTyper_SVs were found in all four lineages (European and North American, wild and domesticated), a surprisingly high proportion given the deep divergence and limited recent gene flow between these groups (Payne et al. 1971; King et al. 2007; Rougemont and Bernatchez 2018) (**Figure 1C**). In contrast, only 17.8% were shared exclusively between wild and domesticated European lineages, and just 0.6% between wild and domesticated North American lineages. This pattern establishes the central question addressed in the following analyses: whether broad SV sharing reflects ancient persistence, recurrent formation at unstable loci, or redistribution through admixture and lineage-specific frequency change. Given the deep divergence and historical isolation between European and North American lineages, these results were not fully in line with expectations: SVs shared across all four lineages were more common than those shared just within lineages of the same continent, despite the very recent split between these domesticated and wild populations. This observation suggests a complex history, involving either long-term maintenance of ancient polymorphisms or highly recurrent SV formation at structurally unstable loci. We revisit these scenarios and their supporting evidence in subsequent sections. In addition, postglacial introgression from European lineages into North America, as well as historical admixture or contamination of domesticated North American stocks with European-origin fish, could also have contributed to the observed sharing (Rougemont and Bernatchez 2018; Wringe et al. 2018; Lehnert et al. 2019; Bradbury et al. 2022; San Román et al. 2025). While these admixture scenarios are not explicitly tested here, they provide alternative, non-mutually exclusive explanations for the extensive SV overlap observed across lineages.

### Identification of large-scale structural variants using PCA-based clustering

Large structural variants (SVs) are known to influence genome evolution at macroscopic scales, contributing to phenotypic divergence, speciation, reproductive isolation, and karyotypic shifts by affecting chromosomal structure, and regulatory landscape (Jin et al. 2023; Ferguson et al. 2024; Bendixsen et al. 2025). However, their detection remains challenging, particularly for variants exceeding the resolution of short-read sequencing (Mahmoud et al. 2019; Gong et al. 2021). To complement our previous SV discovery, we applied a window-based PCA clustering approach (see **Materials and Methods**, Mérot *et al*., 2021), which is designed to detect genomic regions with unusual haplotype structure potentially indicative of large SVs.

Using this method, we identified 25 genomic regions, ranging from 16.9 kb to 3.1 Mb, hereafter referred to as “Lostruct_SVs” (**Supplementary Figure 4, Supplementary Table 1**). These regions were further investigated using population genetic statistics to assess signatures consistent with large SVs. In most cases, heterozygous individuals showed elevated heterozygosity and nucleotide diversity (π), and comparisons between homozygote classes revealed high values of d_XY_ and F_ST_, consistent with strong differentiation and structural divergence (**Figures 3 and 4**). In addition, we observed that all identified SVs were present in at least one copy in all lineages. Because the local PCA approach was not designed to preferentially detect variants shared across all lineages, the presence of all 25 Lostruct_SVs in every lineage was unexpected and motivated further evaluation of their evolutionary origins.

**Figure 3:**
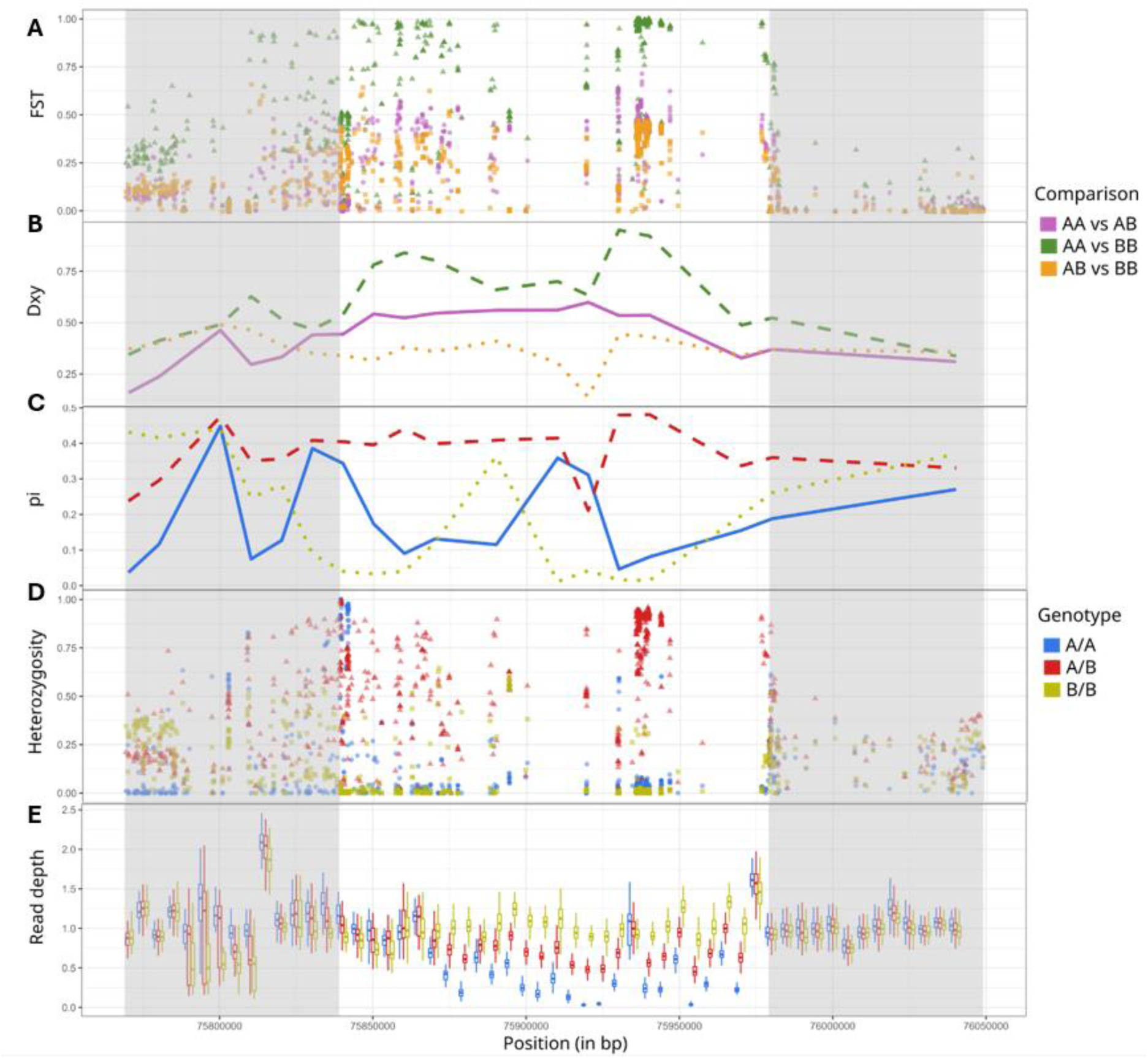
Population genetic summary statistics across one Lostruct_SV region. The SV represented is located on ssa03:75839027 – 75979106. Each panel shows variation in genetic statistics along genomic position (x-axis) for the three inferred genotypes of the SV (A/A, A/B, B/B). Grey shading indicates the boundaries of the putative SV A: FST for each genotype comparison. B: dxy for each genotype comparison. C: Pi for each genotype: D: Heterozygosity for each genotype. E: Read depth for each genotype. The A/A genotype has a near-0 read depth for a large subregion, except for some parts due to collapsed sequences (**Supplementary figure 6**). Alt text: Multi panel figure representing various genomic indicators within a genomic region (lostruct_SV ssa03:75839027 – 75979106) and comparing different genotype or alleles for the SV.

**Figure 4.**
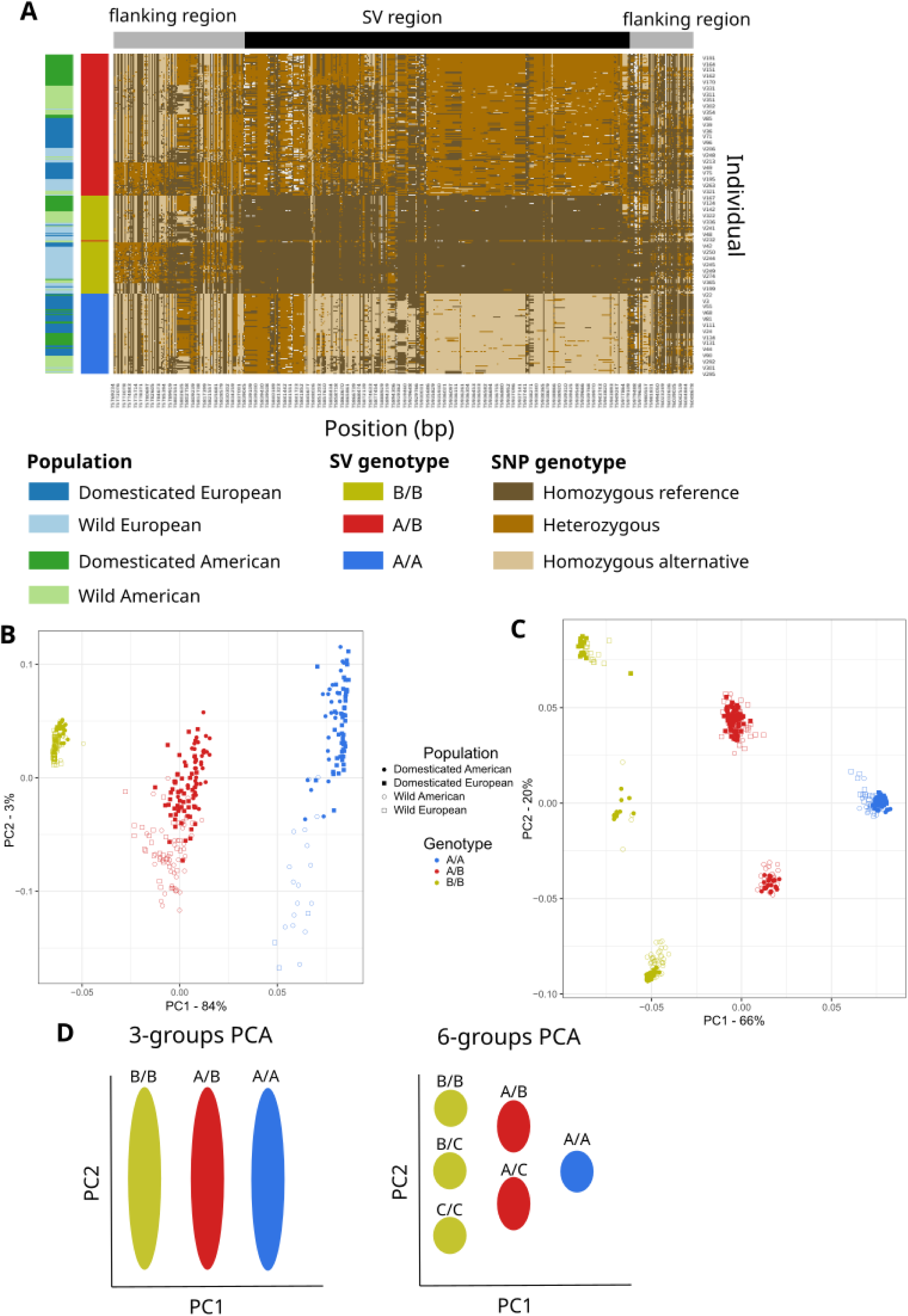
Genotypic clustering and haplotype structure of large structural variants detected by local PCA. **(A)** Haplotype visualization for the lostruct_SV ssa03:75839027 – 75979106. The heatmap represents individual SNP genotypes relative to the reference genome (light = homozygous reference, dark = homozygous alternative, intermediate = heterozygous). Colored bars on the left indicate lineage (green = Domesticated American, light green = wild American, blue = domesticated European, light blue = wild European). Bars on the right indicate the SV genotype inferred by *invClust* (yellow = A/A, red = A/B, blue = B/B). **(B, C)** Principal component analyses (PCA) within two representative SV regions (ssa03:75.8–75.9 Mb **(B)** and ssa05:44.0–44.1 Mb **(C)**). Each point represents an individual. Color denotes SV genotype (yellow = A/A, red = A/B, blue = B/B); shape denotes lineage (filled circle = domesticated American, open circle = wild American, filled square = domesticated European, open square = wild European). **(D)** Schematic representation of PCA cluster configurations among the 25 Lostruct_SVs. Left panel shows the typical three-group pattern expected for a biallelic SV (A/A, A/B, B/B). Right panel shows an unusually six-group pattern, potentially reflecting more complex or recurrent variation. Ellipses illustrate the relative position and separation of genotype clusters. Alt text: A: haplotype visualisation for lostruct_SV ssa03:75839027 – 75979106, showing three distinct haplotype groups; B and C: PCA for two genomic regions (ssa03:75.8–75.9 Mb and ssa05:44.0–44.1) showing grouping of individual according to the SV genotype, with a noticeable six groups clustering in C; D: theoretical PCA representation of genotypes explaining pattern observed in B and C

To further examine whether the regions detected by *lostruct* corresponded to discrete structural variants, we inspected the local haplotype structure within each candidate region. We performed principal component analyses (PCA) based on SNPs located within the putative SV boundaries to visualize potential genotype clusters among individuals. In most regions, the PCA revealed three clearly separated clusters, corresponding to the expected genotypic classes of a biallelic SV (AA, AB, and BB). However, a subset of regions (7 to 14) displayed a more complex pattern, with six partially overlapping clusters (**Figure 4D**). This unexpected pattern suggested the possibility of more than two alleles segregating at these loci, as observed in other species (González et al. 2014; Knief et al. 2016). To evaluate this hypothesis, we compared heterozygosity across the six groups. If three distinct alleles were present, we would expect the three heterozygous combinations to show elevated heterozygosity relative to homozygotes. This pattern was not consistently observed (**Supplementary Figure 5**), providing limited support for a true multiallelic model. Nonetheless, these regions illustrate that haplotype structure in large SVs can be unexpectedly complex and may not always conform to simple biallelic expectations.

To test whether the large SVs detected through haplotype structure reflected underlying copy number variation (CNV), we examined normalized sequencing read depth across all 25 Lostruct_SV regions, stratified by SV genotype. Several regions, including ssa03:75.8–75.9 Mb, ssa03:81.2–81.3 Mb, and ssa27:11.8–12.3 Mb, showed clear genotype-specific depth shifts, consistent with deletions or duplications contributing to the SV signals detected via local PCA. Among these, the region ssa03:75.8–75.9 Mb exhibited one of the most pronounced CNV-associated patterns (**Figure 3E**), consistent with a deletion except for some repeat rich locations (**Supplementary figure 6**). BLAST searches against the *Danio rerio* genome revealed that this region, hereafter termed IG_V_region, was particularly rich in immune-related genes: 8 out of 10 annotated genes showed high similarity to immunoglobulin sequences in other teleost fish species (≥85% identity, E-value < 1e–4) (**Supplementary table 2**). Given the known rapid evolution of immune gene families and the high density of segmental duplications in this region, the structural variation in IG_V_region likely reflects both genomic instability and immune-related functional specialization. Overall, no significant Gene Ontology (GO) enrichment was detected (FDR < 0.05) on the 21 Lostruct_SV regions overlapping annotated genes and many regions contained poorly annotated genes.

### Contrasting frequency patterns of large SVs within and between continents

To assess how large structural variants (SVs) are retained across lineages, we examined allele frequencies of the 25 Lostruct_SVs identified through haplotype-based clustering. These SVs were prioritized because they show clear haplotype differentiation signals and, due to their large size and proximity to functional regions, are likely to have stronger effects on linked variation than smaller SVs.

Within each continent, we observed a strong positive correlation in SV frequencies between wild and domesticated lineages (Europe: slope = 0.65, R² = 0.4, p = 7.3 x 10⁻⁴; America: slope = 0.83, R² = 0.79, p = 3 x 10⁻^9^; **Figure 5A,B**). This pattern is consistent with expectations given the relatively recent origin of domestication (∼15 generations). A single outlier, IG_V_region, the immune-related region on chromosome 3, showed a pronounced frequency increase in domesticated European salmon (0.69) relative to wild counterparts (0.19), deviating beyond the 95% confidence limits of the regression line. This deviation could result from either a founder effect, domestication-associated selection, or both.

**Figure 5.**
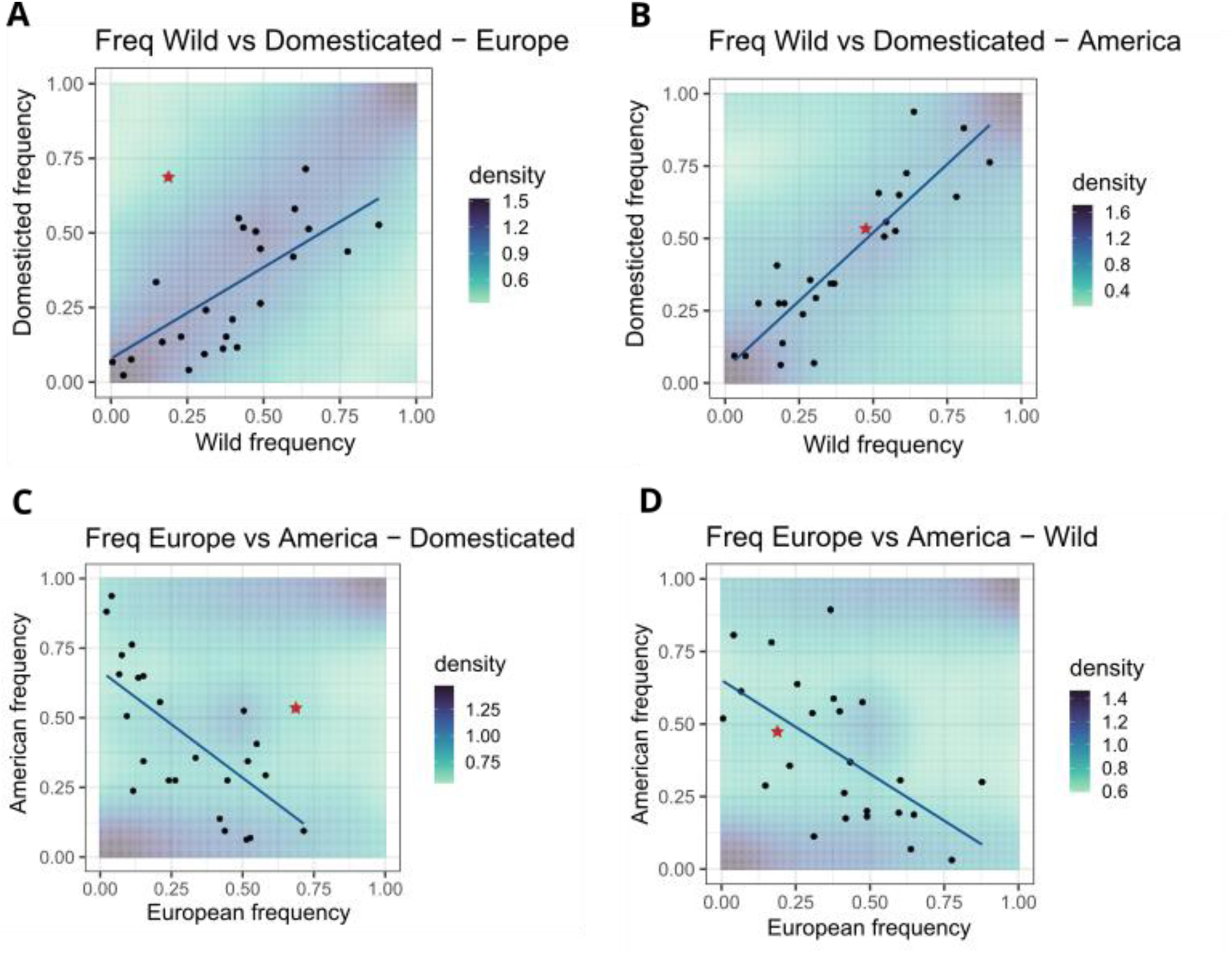
Frequency correlations of large structural variants (Lostruct_SVs) among populations. (A, **B)** Allele frequencies of 25 Lostruct_SVs in wild (x-axis) versus domesticated (y-axis) populations for Europe **(A)** and North America **(B)**. Each point represents a single SV; background shading indicates the frequency density of bayestyper_SVs. The red star marks IG_V_region, which shows a strong frequency shift in domesticated Europe. **(C, D)** Comparison of SV frequencies between European (x) and American (y) populations for domesticated **(C)** and wild **(D)** groups. Blue lines show regression fits; shading indicates the frequency density of bayestyper_SVs. The red star again corresponds to IG_V_region. Alt text: 4 panels representing different lineage-lineage comparison of Lostruct_SVs alleles frequencies, with colored background representing Bayestyper_SVs alleles frequencies. Comparison within continent (for instance wild vs domesticated frequency within European lineages) display a positive correlation, while comparison within domestication status (for instance European vs American frequency within domesticated lineages) display a negative correlation.

When comparing European and North American lineages, we expected a weakly positive correlation because many SVs likely predate continental divergence. Contrary to this expectation, allele frequencies were negatively correlated in both domesticated (slope = –0.77, R² = 0.43, p = 3.6 × 10⁻⁴; **Figure 5C**) and wild (slope = –0.64, R² = 0.36, p = 1.4 × 10⁻⁴; **Figure 5D**) groups. This contrast with the within-continent pattern suggests that ancient SVs may have followed different frequency trajectories across continents, potentially due to geographically variable selection, demographic history, or linked effects.

Previous SNP-based selection scans, including XPEHH analyses conducted in the same American and European lineages pairs investigated here, reported several candidate regions under artificial selection during domestication (Buso et al. 2025). Among the SVs identified in the present study, 3 Lostruct_SVs in the American population and 3 SVs (1 Lostruct_SV, 2 Bayestyper_SVs) in the European lineage overlapped with these previously reported candidate regions (**Supplementary Figure 7, Supplementary Table 3)** (FDR = 10^-4^). This overlap suggests that some SVs occur in regions previously identified as candidates for domestication-associated differentiation, although the signal may reflect selection, reduced recombination, or linked variation around the SV.

### Simulated evolutionary scenarios reveal distinct retention patterns of ancient and recurrent SVs

To better understand the evolutionary processes shaping SV diversity, we simulated forward-time models of SV evolution under different scenarios using SLiM v4.2.2 (Haller and Messer 2023) (**Materials and Methods**). This analysis was motivated by the unexpectedly high proportion of SVs shared across the four sampled lineages, which could result either from recurrent mutations or from long-term maintenance of ancient polymorphisms by selection.

Ancient SVs (SVs arising before lineage divergence) were consistently more likely to be retained across all lineages (99% and 40%) than recurrent SVs (17% and 15%) (**Figure 6A**), regardless of the recurrence rate (**Supplementary Figure 8**). Under balancing selection, recurrent SVs persisted in more lineages than under directional selection (2.42 lineages on average vs 2.29). In contrast, ancient SVs were more likely to persist in more lineages under directional selection than under balancing selection (4 lineages on average vs 3.27). Ancient SVs with directional selection tended to become rapidly fixed and were therefore, excluded from further analysis, as fixed variants would not be detectable in lineage comparisons (**Figure 6A**). Overall, the simulations indicate that SVs shared across all four lineages are more likely to represent ancient polymorphisms than recurrent events.

**Figure 6:**
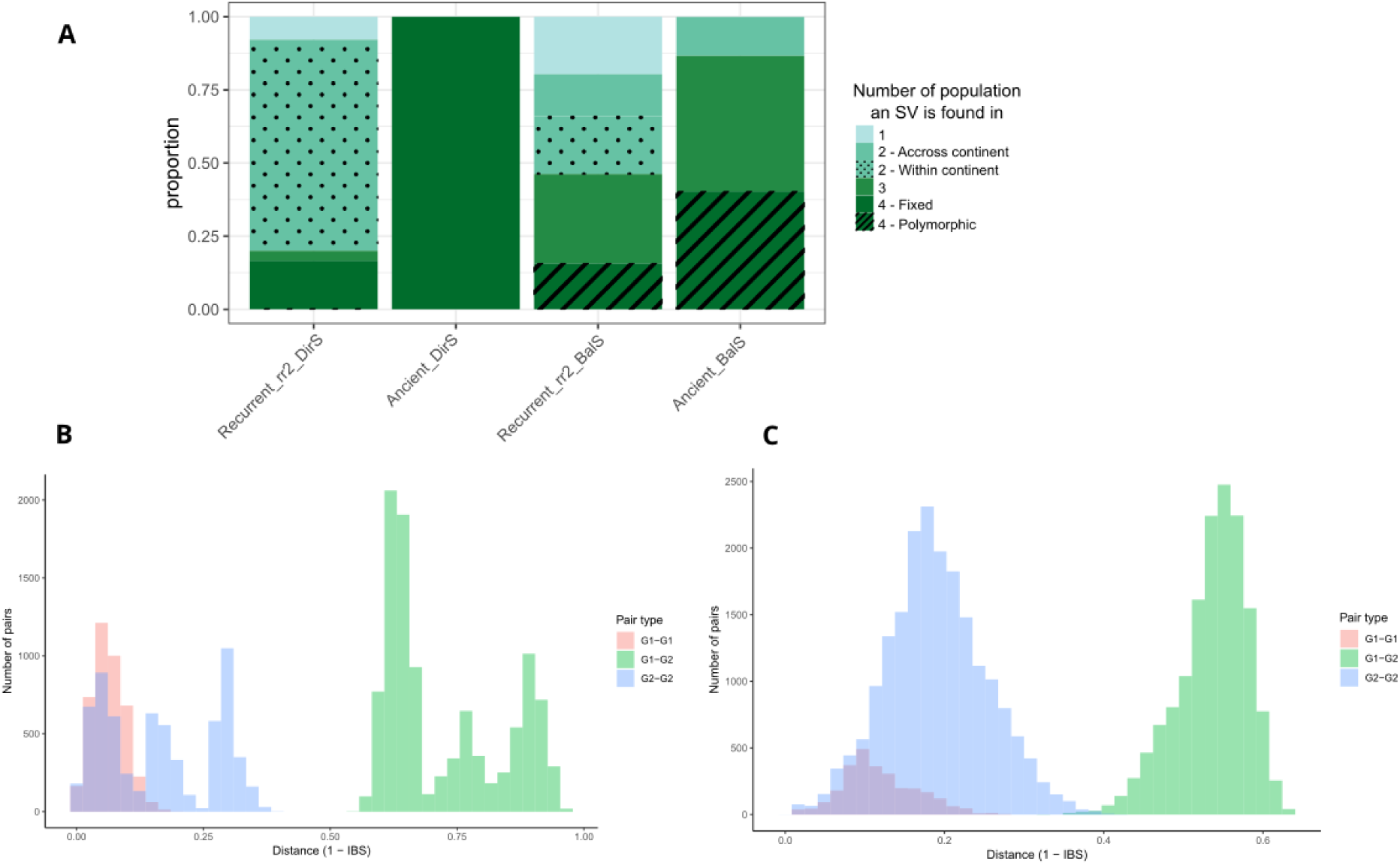
Evolutionary history of simulated SV and inferred recurrent SVs. **A:** Labels indicate the history of SVs (recurrent or ancient), as well as the selection type (directional selection = DirS; balancing selection = BalS) and the recurrent rate (rr2 = 2 SVs per simulation on average). Simulations were run with a selection coefficient of 1.1. **B-C**: Genetic distance between SNPs in haplotype, G1-G1: AA vs AA; G1-G2:AA vs BB; G2-G2: BB vs BB, for two lostruct_SVs. Under recurrent SV, the distribution is expected to be multimodal for G1-G2 (**B**, ssa05:44020000-44084228), which is not the case for other SVs (**C**, ssa12:63150000-63800000) Alt text: A: Stacked bar plot indicating the number of lineages a Sv was retained in after the simulation, for the four different scenarios. B and C: Histograms representing the genetic distance between haplotypes for two lostructs_SVs, displaying a mutimodal distribution for G1-G2.

To further characterize the evolutionary origins of the detected structural variants, we examined the distribution of pairwise haplotype distances (1 − IBS) among individuals homozygous for alternative SV genotypes. For each SV, we compared three distance categories: between individuals homozygous for haplotype 1 (G1–G1, AA vs AA), between individuals homozygous for haplotype 2 (G2–G2, BB vs BB), and between the two homozygote groups (G1–G2, AA vs BB). Forward simulations indicated that SVs of ancient origin typically yield unimodal distance distributions, with within-group distances forming a single peak and between-group distances showing a single, broader mode corresponding to deep divergence between haplotypes. In contrast, recurrently arising SVs are expected to produce multimodal distributions, as independent mutational events generate several distinct haplotypes within the same genotypic class.

When applying this simulation-informed framework to the empirical data, most SVs showed the unimodal “ancient” pattern. However, four loci exhibited multimodal distance distributions consistent with recurrent origin. A representative example is shown in **Figure 6B**. In this case, distances among G1 individuals (G1–G1, AA vs AA, red) form a single narrow peak, indicating a uniform haplotype background. In contrast, distances among G2 individuals (G2–G2, BB vs BB, blue) and between G1 and G2 (G1–G2, AA vs BB, green) form three distinct peaks, reflecting the coexistence of multiple divergent haplotypes classified under the same genotypic state. This pattern implies that the G2 allele has arisen independently more than once, producing several haplotype lineages that now segregate in the population. All four loci displaying this pattern (ssa16:33750000-34020915, ssa05:44020000-44084228, ssa16:60620000-60880000, ssa21:38333640-38459102, **Supplementary Figure 9**) also exhibited complex six-cluster structures in PCA space (exemplified in **Figure 4D**). We found that these four regions were surrounded by more TEs than random regions but not significantly more than other Lostructs_SVs (**Supplementary Figure 10**). For BayesTyper_SVs, similar analysis did not reveal clear evidence of recurrent SVs, in particular due to the short size of these SVs not being well suited for this haplotype comparison method. Thus, broad sharing across all lineages and recurrent origin represent distinguishable, rather than interchangeable, explanations for shared SVs.

## Discussion

Structural variants shared among populations are often interpreted as evidence of shared ancestry, gene flow, or recurrent mutation, but these alternatives can produce similar presence–absence patterns and are difficult to distinguish from sharing alone.Here, we combined short-read SV genotyping, haplotype-based clustering, and forward simulations to investigate the evolutionary history of SVs across four Atlantic salmon lineages differing in geographic origin and domestication status. This integrative framework allowed us to contrast ancient polymorphisms maintained across lineages with recurrent mutations occurring at structurally variable genomic regions.

Nearly half of the SVs were shared among all four lineages, suggesting the long-term persistence of ancestral polymorphisms, while a small number of large SVs (7 to 14) exhibited multimodal haplotype-distance distributions and complex six-cluster PCA patterns indicative of recurrent origin. One notable SV on chromosome 3 (IG_V_region), encompassing immunoglobulin-related genes, showed a marked frequency increase in domesticated European salmon, consistent with a domestication-associated frequency shift. These findings reveal that the SV landscape of Atlantic salmon reflects both deep evolutionary maintenance and ongoing structural turnover.

Structural variants in Atlantic salmon were strongly enriched within or near repetitive regions, particularly segmental duplications and LTR retrotransposons, indicating that the genomic context plays a major role in their formation. Such enrichment is consistent with the well-established role of repetitive sequences in promoting non- allelic homologous recombination (NAHR) and replication-based rearrangements (Kidd et al. 2010; Carvalho and Lupski 2016). Segmental duplications provide extensive sequence homology that facilitates ectopic pairing, leading to recurrent deletions, duplications, or inversions at the same loci (Emanuel and Shaikh 2001). The strong enrichment we observed among LTR retrotransposons further suggests that retroelement proliferation contributes to ongoing genome instability. Because LTR elements propagate by a copy-and-paste mechanism, they can mediate unequal crossover and generate large insertion–deletion polymorphisms, whereas DNA transposons, which rely on a cut-and-paste mechanism, are less likely to promote large rearrangements (Gray 2000; Burssed et al. 2022).

These observations indicate that SV formation in the Atlantic salmon genome is not random but is biased toward structurally variable regions. This bias provides a mechanistic explanation for why certain loci repeatedly generate the same or hihgly similar SVs, as seen for the recurrent variants detected in later analyses. The co-localization of SVs with repetitive and duplicated regions thus highlights the role of intrinsic genome architecture in shaping the mutational landscape of large structural variation.

### Shared SVs across lineages and the persistence of ancestral polymorphisms

Nearly half of the high-confidence Bayestyper_SVs were shared among all four Atlantic salmon lineages, an unexpectedly high proportion given the deep divergence and long-term geographic isolation between European and North American lineages (Payne et al. 1971; Rougemont and Bernatchez 2018). This extensive sharing cannot be readily explained by recurrent mutation alone, as our simulations showed that recurrent SVs are rarely retained across multiple lineages. Instead, it indicates that a large fraction of SVs represents ancient polymorphisms that have persisted since before the continental split.

The preferential localization of these shared SVs in intronic and intergenic regions further supports this interpretation, as such regions are less affected by purifying selection and more likely to tolerate long-term structural variation. Similar patterns have been reported in other taxa, where chromosomal inversions or copy-number variants have been maintained for millions of years under balancing or fluctuating selection (Lindtke et al. 2017; Göktay et al. 2021; Saitou et al. 2021; Aqil et al. 2023). The persistence of ancestral SVs may, therefore, reflect a combination of neutrality and balancing forces that prevent fixation or loss. Moreover, within each continent, SV frequencies were positively correlated between wild and domesticated lineages, consistent with recent shared ancestry and limited time for frequency divergence.

However, post-glacial secondary contact and introgression between European and North American lineages could have contributed to the persistence and redistribution of some SVs (Rougemont and Bernatchez 2018; Lehnert et al. 2019). Therefore, while many shared SVs are consistent with ancient polymorphisms predating the continental split, secondary contact may also have shaped their present-day distribution.

### Large-scale SVs and complex haplotype structures

While most SVs exhibited patterns consistent with long-term maintenance, a subset of large-scale variants revealed unexpectedly complex haplotype structures. Using local PCA analyses, we identified 25 genomic regions (Lostruct_SVs) showing pronounced clustering of haplotypes, consistent with the presence of large rearrangements. In most regions, individuals grouped into three discrete clusters corresponding to the genotypic classes expected for a biallelic SV (AA, AB, and BB). However, several loci deviated from this pattern and displayed six partially overlapping clusters, implying that multiple distinct haplotypes segregate at the same genomic position. Previous studies have reported similar multilayered clustering patterns, reflecting complex structural variants and, in some cases, providing evidence for tri-allelic SV loci. (González et al. 2014; Knief et al. 2016; Lehnert et al. 2025). However, some of the SVs identified in our study did not align with a simple tri-allelic model. In particular, the heterozygosity patterns and haplotype-distance distributions did not match the expectations of three stable allelic classes segregating within the population. Instead, the clustering patterns suggested that multiple structurally similar variants had arisen on distinct haplotype backgrounds.

Our forward simulations predicted that recurrent SVs could produce this type of multi-cluster haplotype configuration. Consistent with this expectation, these loci exhibited multimodal haplotype-distance distributions in the empirical data, indicating the presence of multiple divergent haplotypes within the same genotypic category. Such recurrent structural changes generate “cryptic” haplotype diversity that may be overlooked when SVs are interpreted as simple, single-origin, biallelic variants, even with long read sequencing methods (Jenko Bizjan et al. 2020; Qin and Li 2025; Schloissnig et al. 2025).

### Functional relevance and domestication-associated frequency change

Among the large structural variants identified, the region IG_V_region on chromosome 3 showed a pronounced allele-frequency increase in domesticated European salmon (0.19 in wild versus 0.69 in domesticated individuals), while no comparable shift occurred in North American populations. Although a founder effect associated with the establishment of domesticated stocks could contribute to the observed frequency shift, the pattern remains consistent with a domestication-associated frequency shift at a structurally unstable immune region, although founder effects and selection cannot be distinguished with the present data. The biological context of the region points to a plausible target of selection. The locus overlaps with a copy-number variable segment enriched for immunoglobulin-like genes, situated within a cluster of segmental duplications. Such architecture is well known to facilitate both recurrent rearrangements and rapid immune-gene evolution (Traherne et al. 2010; Lin and Gokcumen 2019; Rodriguez et al. 2023). Read-depth profiles further indicate that this locus shows genotype-specific shifts in copy number, consistent with both copy-number gains and losses across the inferred SV genotypic classes. Under aquaculture conditions, characterized by high population density and elevated pathogen exposure, selection on immune-related copy-number variation is plausible (Krkosek 2010; Li et al. 2017; Buso et al. 2025).

Similar immune-related SVs have been implicated in adaptation to parasite pressure and disease resistance in other species, supporting the notion that structural variation can serve as an important substrate for rapid immune evolution (Penso-Dolfin et al. 2020; Göktay et al. 2021; Karageorgiou et al. 2024). The example of IG_V_region thus illustrates how functional gene content and artificial selection can shape the evolutionary trajectory of specific SVs in domesticated salmon.

### Frequency divergence across continents

The contrasting patterns of allele-frequency correlations within and between continents inform the evolutionary interpretation of large SVs in Atlantic salmon. Within each continent, allele frequencies were positively correlated between wild and domesticated lineages, consistent with recent shared ancestry and limited divergence since domestication. In contrast, allele frequencies were negatively correlated between European and North American populations in both wild and domesticated groups. Recurrent mutation alone is unlikely to account for this pattern, as it would not be expected to produce consistent inverse frequency shifts across multiple loci. A more plausible explanation is that many of these SVs predate continental divergence and have subsequently followed different frequency trajectories in each region. This divergence may reflect direct effects of the SVs themselves, selection on linked variants, demographic history, or other lineage-specific processes in different environmental conditions, including temperature regimes, photoperiod, and pathogen communities (Lehnert et al. 2020). Notably, previous work has reported cases in which associations between large haplotypes and life-history traits differ in direction between European and North American lineages, indicating that similar genomic regions can be linked to contrasting phenotypic outcomes across continents (Lehnert et al. 2020). Over extended timescales, reduced recombination or haplotype structure around SVs could further expand these continent-specific differences.

Environmental contrasts may not be strictly comparable across the studied populations on either side of the Atlantic. For example, oceanographic differences, such as the influence of the Labrador Current along the Canadian Atlantic coast, result in markedly colder marine environments at comparable latitudes relative to Northern Europe (Drinkwater et al. 2013). Thus, even with similar fitness landscapes, local habitat differences could result in selection for different allele frequencies between continents, potentially contributing to the observed inverse frequency patterns. Alternatively, temporally fluctuating selection, in which the relative advantage of alternative alleles changes through time, could also generate continent-specific frequency differences (Lehnert et al. 2020), although that require the environments to be consistently in different phases and so selection on SV genotypes to be out of phase between the continents. Together, these scenarios provide plausible mechanisms by which ancient SVs may be retained while exhibiting contrasting frequency trajectories across continents.

The observed negative correlation was only observed for large SVs (Lostruct_SVs), raising questions about the differences between short and large SVs leading to different observed patterns. One possibility is that large SVs are more prone to accumulate mutations than short ones, mutations which could be the proximal effect on fitness (Weischenfeldt et al. 2013). Another explanation resides in the fact that large SVs are more likely to affect several genes, which can easily lead to a greater allele complexity than shorts SVs, with more complex selection patterns.

These findings imply that the SV landscape of Atlantic salmon reflects a long-term balance between geographically variable selection, and historical demographic events. Ancient SVs have persisted across continents, but their allele frequencies have been reshaped by region-specific evolutionary processes, resulting in parallel yet directionally opposed genetic landscapes for larger SVs.

### Simulations and inference of recurrent SVs

Forward-time simulations provided a theoretical context for interpreting how different evolutionary processes can produce the observed patterns of SV sharing. Under a wide range of parameters, recurrent mutations were predicted to be short-lived or geographically restricted, whereas ancient variants were expected to persist across lineages under neutrality or balancing selection. These simulations indicate that widespread SVs shared across lineages are more consistent with long-standing polymorphisms than recurrent mutation alone.

Applying this simulation-informed framework to empirical data allowed us to identify a small number of loci consistent with the recurrent scenario. Four Lostruct_SVs exhibited multimodal haplotype-distance distributions (1–IBS) and complex six-cluster PCA configurations, patterns predicted by simulations for recurrent mutations. In these loci, one haplotype class was genetically homogeneous, whereas the alternative class comprised multiple divergent haplotypes, consistent with similar structural rearrangements having arisen independently at the same genomic sites. Such patterns are difficult to explain by ancestral polymorphism alone and are instead consistent with recurrent mutation in regions of structural instability. The convergence of structural and population-genetic evidence supports recurrent SV formation as an ongoing process in the Atlantic salmon genome. Although these events are relatively rare compared to the abundance of ancestral SVs, their occurrence highlights the dynamic nature of genome evolution, where ancient variation provides continuity while recurrent mutation introduces new genomic configurations.

### Methodological considerations and future directions

Our approach combines population-scale short-read genotyping, haplotype-based clustering, and forward simulations to investigate the evolutionary history of structural variants. While this integrative framework provides a coherent view of SV dynamics, several technical aspects warrant consideration for future refinement. Short-read sequencing inevitably limits the resolution of complex rearrangements, particularly at repetitive breakpoints, where mapping ambiguity can lead to merged or fragmented variant calls. Because our pipeline was conservative against false positives and ambiguous variants, some true variants were likely missed; however, the variants retained after stringent filtering are expected to represent a highly reliable subset of the SV landscape.

The identification of recurrent-like SVs was based on haplotype-distance patterns derived from simulations and qualitative inspection of multimodality. Although not strictly quantitative, this simulation-informed criterion provides a consistent and transparent basis for distinguishing recurrent from ancient origins. Similar qualitative approaches have been successfully applied in studies of inversion polymorphisms and complex haplotype architectures in natural populations (Boettger et al. 2016; Xu et al. 2017; Porubsky et al. 2020; Schaal et al. 2022). Future work integrating long-read sequencing, optical mapping, and direct breakpoint reconstruction will allow more precise quantification of recurrence rates and mutational mechanisms.

As an additional consideration, hybridization was not explicitly modeled in our analytical framework. Although the lineages examined here are generally thought to have experienced long-term geographic isolation, postglacial introgression events have been documented in parts of their range, and traces of European ancestry may persist in some domesticated North American populations (Bradbury et al. 2022). Such histories, if present, could contribute to a small fraction of shared SVs. Future analyses incorporating explicit admixture models will help clarify the extent to which hybridization has influenced the observed patterns.

Finally, integrating structural variation with transcriptomic and phenotypic data will be essential to evaluate the functional consequences of SV polymorphisms. Across diverse taxa, SVs have been shown to contribute to gene expression variation, regulatory divergence, and adaptive phenotypes beyond what is captured by SNP-based analyses alone (GTEx Consortium et al. 2017; Alonge et al. 2020; Scott et al. 2021; Falker-Gieske et al. 2022; Zhang et al. 2024). In Atlantic salmon, applying comparable integrative approaches allows these general effects to be examined in a system shaped by recent and rapid domestication, deep continental divergence, and a structurally dynamic genome influenced by whole-genome duplication and lineage-specific chromosomal rearrangements.

In conclusion, our results show that shared SVs across divergent lineages cannot be treated as a single evolutionary class. In Atlantic salmon, broad SV sharing is largely consistent with ancient persistence, whereas a smaller subset of large SVs shows patterns consistent with recurrent formation, and some structurally complex regions exhibit lineage-specific frequency shifts. By explicitly contrasting alternative origins of shared SVs, this study provides a framework for interpreting structural variation across natural populations and evolutionary timescales.

## Material and methods

### Sampling design and whole-genome sequencing data

We analyzed whole-genome sequencing data from four Atlantic salmon lineages representing two continents (Europe and North America) and two domestication status (wild and domesticated). In total, 369 individuals were initially considered. Farmed North American samples (n = 80) originated from three source rivers in North America (NCBI Project PRJEB34225). Farmed European samples (n = 112) belonged to the MOWI strain, derived from the River Bolstad in the Vosso drainage, the River Årøy, and marine captures near Osterfjord and Sotra in Western Norway (Glover *et al*., 2009, NCBI Project No. PRJEB47441). As wild counterparts, we used 79 wild North American individuals collected from eight rivers in Québec and 98 wild European individuals from 17 rivers in Western Norway (Bertolotti *et al*., 2020, NCBI Project No. PRJEB38061).

Paired end Illumina reads from all individuals were mapped to the Atlantic salmon reference genome Ssal_v3.1 (GCA_905237065.2) using bwa-mem2 (Li and Durbin 2009). One wild North American sample that could not be reliably genotyped for structural variants was removed from downstream analyses.

Two Atlantic salmon individuals, derived from a breeding nucleus of an aquaculture company (MOWI) (NCBI Project PRJEB113180), were used for long-read sequencing. These individual were used to investigate coverage of the IG_V_region (**Supplementary figure 6**).

### SNP detection

SNP calling followed the pipeline described in Buso *et al*., 2025 for the mapping and variant calling. Mapping, variant calling and quality control were performed independently for each of the four datasets (Farmed Norwegian, Wild Norwegian, Farmed North American, Wild North American) using the same pipeline and parameters. Illumina reads were mapped to the Atlantic salmon genome (Ssal_v3.1; GCA_905237065.2) using bwa-mem2 (Li and Durbin 2009). Duplicate reads were marked with gatk4-spark:4.3.0.0 MarkDuplicates. Genetic variation was identified using gatk4-spark:4.3.0.0 HaplotypeCaller with default parameters and the individual genotypes were merged with gatk4-spark:4.3.0.0 GenotypeGVCFs. To avoid the detection of false positives, variants were filtered by the following parameters with gatk4-spark:4.3.0.0 (McKenna et al. 2010) : “QD < 2.0” which filters out variants with a low quality score by depth of coverage; “QUAL < 50.0” to sort variants with a low-quality score; SOR > 4.0 that removes variants where the number of reads supporting the reference allele is significantly skewed towards one strand (forward or reverse) compared to the other; FS > 60.0: to filter out variants whose reads supporting the reference allele versus the alternative allele are significantly different between the two strands; and finally ReadPosRankSum < -8.0: to remove variants with a low score for the difference between the mean position of the reference allele and the alternative allele among the reads supporting the variant (https://gatk.broadinstitute.org/hc/en-us).

Variants detected in all lineages were then merged together in a single file using bcftools v.1.9 (Danecek et al. 2021), in order to homogenize variants across all populations. Variants with missingness higher than 10% were removed using vcftools v0.1.16 (Danecek et al. 2011).

### Read-based structural variant discovery and genotyping

Initial discovery of structural variants (SVs) relied on read-based calling. For each individual, we used Manta v1.6.2 (Chen et al. 2016) to detect candidate SVs using paired-end and split-read information. Within each lineage, individual-level calls were merged using SURVIVOR v1.0.7 (Jeffares et al. 2017) with parameter “100 1 0 0 0 30”. in order to cluster SVs with similar breakpoints and to construct a lineage-wide candidate SV set. Variants were first filtered by Manta quality score, retaining only SVs with QUAL > 100 (**Supplementary Figure 1**).

Because the Manta pipeline alone does not clearly distinguish between truly absent variants and uncalled genotypes, and because precise genotyping was essential for comparative analyses, we refined SV genotypes using BayesTyper v1.5 (The Danish Pan-Genome Consortium et al. 2018) BayesTyper uses a k-mer based probabilistic model that combines read support and expected genotype configurations. We provided BayesTyper with the SVs discovered by Manta and followed the batch processing recommendations from the official documentation (**Supplementary Figure 1**). Due to computational limitations, individuals were divided into batches of five, yielding 74 batches that were genotyped separately and then merged using bcftools (Danecek et al. 2021) as recommended in the bayestyper guidelines.

We removed SVs with more than 75 % missing genotypes and a BayesTyper quality score below 10 using vcftools v0.1.16 (Danecek et al. 2011). During the BayesTyper step, information on SV type was not retained, so SV types were reassigned by matching the genomic coordinates of BayesTyper-genotyped variants to the original Manta calls (**Supplementary Figure 1**). We identified one individual that appeared as a consistent outlier in genome-wide principal component analyses was excluded from population genetic analyses using this data set. The final high-confidence SV set, referred to as BayesTyper_SVs, contained 6,683 variants and formed the basis for genome-wide analyses of SV size distribution, genomic context, and sharing across lineages.

### Annotation of genomic context and repetitive elements

Gene annotations for the Ssal_v3.1 reference genome were used to assign SVs to genomic features. We used bedtools v2.30 (Quinlan and Hall 2010) to intersect BayesTyper_SVs with gene models and to classify variants according to their location in exons, introns, or intergenic regions.

To investigate the association between SVs and repetitive elements, we first built a de novo transposable element (TE) library for Atlantic salmon using RepeatModeler v2.0.4 (Flynn et al. 2020). Elements classified as “unknown” by RepeatModeler were reclassified using DeepTE tool (Yan et al. 2020). To reduce redundancy among TE families and to avoid double counting similar families under different names, we clustered the TE consensus sequences using CD- HIT v4.8.1 following recommended parameters from published TE curation workflows (Goubert et al. 2022).

Genome-wide TE annotation was then performed using the curated TE library. Segmental duplication tracks were retrieved from the UCSC Genome Browser for Ssal_v3.1. Using bedtools v2.30 (Quinlan and Hall 2010), we calculated the proportion of BayesTyper_SVs that overlapped or were located within 5 kb of each repeat category, including segmental duplications, LTR retrotransposons, DNA transposons, and other TE classes. To assess enrichment relative to random expectation, we generated null distributions by simulating sets of random genomic intervals matched to the observed SVs in size and chromosome and computing the same overlap statistics.

To investigate the Tc1 mariner peak observed in the SV size distribution (Bertolotti et al. 2020), we extracted sequences for SVs in the 1,432-1,436 bp size range and aligned them against the Tc1 mariner reference sequence EF685967.1 using BLASTn (Korf et al. 2003). SVs showing at least 95 % sequence identity were classified as Tc1 mariner associated.

### Detection of large structural variants using local PCA

Large structural variants are difficult to detect via short-read based callers which often can not span the whole SV. To complement the read-based pipeline, we used a local SNP-based PCA approach to detect large SVs implemented in the R package *lostruct* (Li and Ralph 2019). Genome-wide SNP data were partitioned into windows of 1,000 SNPs along each chromosome, and principal component analyses were performed in each window using the run_lostruct.R script. We then used the built-in “summarize_run.Rmd” script to identify windows with extreme PCA configurations relative to the genomic background.

Most windows displayed PCA patterns typical of genome-wide background structure and were therefore not retained for further analysis. Instead, we focused on windows exhibiting clear deviations from this background pattern, which are indicative of structurally divergent haplotypes expected under the presence of large SVs. These atypical windows were subsequently clustered using k-means based on their major principal components. For each candidate region, we merged adjacent windows with similar PCA structure, resulting in 51 putative SV regions.

For these regions, we calculated F_ST_ between the inferred genotype groups using vcftools and visually inspected F_ST_ along the chromosome. Regions showing extended blocks of elevated F_ST_ were retained, and their approximate breakpoints were refined based on the F_ST_ profiles. Genotypes for each region were then assigned using *invClust* (https://github.com/isglobal-brge/invClust/), which infers karyotypes of inversion-like polymorphisms from SNP data. Regions in which only two genotype classes could be confidently resolved were excluded. After this filtering, 25 regions remained and are referred to as Lostruct_SVs (**Supplementary figure 5**, **Supplementary table 1)**.

### Population genetic statistics and potential copy number variants

For each Lostruct_SV, we calculated population genetic statistics within and between inferred genotype classes. Heterozygosity and nucleotide diversity (π) were computed within each genotype using the R packages *vcfR* and *adegenet* (Jombart 2008; Knaus and Grünwald 2017). Between-genotype differentiation was summarized using dXY and FST, calculated with pixy v1.2.10.beta2 and vcftools respectively (Danecek et al. 2011; Korunes and Samuk 2021).

To test whether Lostruct_SVs corresponded to underlying copy number variation, we estimated per-base read depth using mosdepth and summarized normalized coverage across SV regions for each genotype class. Regions showing clear genotype-specific coverage shifts were interpreted as candidates for deletions or duplications contributing to the haplotype structure detected by local PCA.

These statistics were computed for the SV region including flanking regions corresponding to 50% of the size of the SVs (upstream and downstream). The PCAs displayed in **Figure 4B and C** were only computed on the SV region itself, to only reflect the structure of the SV.

### Gene content and functional annotation of large SVs

For Lostruct_SVs overlapping annotated genes, we extracted gene models using bedtools and summarized gene content. To investigate the gene content of the immune-related region (IG_V_region), we conducted BLASTn (Korf et al. 2003) searches of its annotated genes against the Danio rerio genome GRCz11 and recorded matches with at least 85 % identity and E-values < 1e-4, which were predominantly immunoglobulin genes.

We used ShinyGO (V 0.80, http://bioinformatics.sdstate.edu/go/) to perform Gene Ontology (GO) enrichment analyses for Lostruct_SVs containing annotated genes. No GO terms remained significant after multiple testing correction at FDR < 0.05.

### Simulations of structural variant evolution

To explore the processes underlying the sharing of SVs among lineages, we performed forward-time simulations using SLiM v4.2.2 (Haller and Messer 2023) following a framework similar to Berdan *et al*., 2021, with different types of selection (directional selection or heterozygote advantage).and different types of evolutionary history (variant being ancestral or appearing in parallel). We modeled a four-population system representing European and North American, wild and domesticated lineages, and simulated structural variants under alternative scenarios.

In “ancient SV” scenarios, SVs arose before lineage divergence and could be maintained or lost in each descendant lineage. In “recurrent SV” scenarios, SVs could arise independently on different haplotype backgrounds after lineage divergence, allowing for multiple mutational origins. For both ancient and recurrent SVs, we considered different selection regimes, including directional selection and balancing selection. For each combination of parameters, we performed 10,000 replicate simulations and retained only those in which the SV remained polymorphic in at least one lineage, because fixed variants would not be detectable in our lineage comparisons.

SNP mutations were simulated with a rate of 1.06 × 10⁻⁸ per site per generation (Stenløkk et al. 2022). Each scenario was run for 2,000 generations. For analysis of haplotype structure, we sampled 50 individual genomes per population from the final generation. Full details of demographic settings, recombination rates, selection coefficients, and recurrence rates are provided as scripts on GitHub.

### Haplotype distance and IBS analyses

To distinguish between ancient and recurrent SV origins, we examined pairwise haplotype distances within and between SV genotype classes. For each Lostruct_SV and BayesTyper_SV, we extracted SNPs within the putative SV boundaries and computed 1 − IBS using PLINK v2 and 1.9 (Chang et al. 2015) treating 1 − IBS as a measure of haplotype divergence.

For each SV, we defined three distance categories, namely distances between individuals homozygous for genotype 1 (G1-G1, AA vs AA), between individuals homozygous for genotype 2 (G2-G2, BB vs BB), and between the two homozygote groups (G1-G2, AA vs BB).

The distributions of these three-distance classes were compared to expectations from the SLiM simulations. Ancient SVs are expected to produce unimodal distance distributions with a single broad mode for between-genotype distances, whereas recurrent SVs are expected to generate multimodal patterns reflecting multiple independent haplotype backgrounds.

We used these patterns to classify candidate recurrent SVs among Lostruct_SVs and to confirm that BayesTyper_SVs, which are typically shorter, did not provide sufficient haplotype information to robustly detect recurrent origins.

## Data and Code availability

All scripts used for read mapping, variant calling, structural variant genotyping, annotation, generation of transposable elements library, population genetic analyses, simulations, and figure generation are available at https://github.com/celiand/SV_evolutionary_history, together with configuration files and example commands enabling full reproducibility of the analyses described in this study.

## Author contribution

The study was conceived by CD, MS and NJB. Data was generated by CD, MA, JSK, MS. Bioinformatic and computational analyses were conducted by CD and MS and DM. Data curation was done by AVPL. Interpretation of the results were done by all authors. Drafting paper was done by CD, MS and NJB. Manuscript editing was done by CD, MS and NJB.

## Acknowledgments

We acknowledge the use of the Orion computing cluster at the Norwegian University of Life Sciences (NMBU). We thank Erik Sandertun Røed from Centre for Ecological and Evolutionary Synthesis – University of Oslo, for his help and advice with SLiM simulations. The study was supported by the Research Council of Norway (grant nos. 325874, 275310, and 221734).

