## Supplementary materials for "Ancient persistence and recurrent emergence of structural variants across divergent Atlantic salmon lineages"

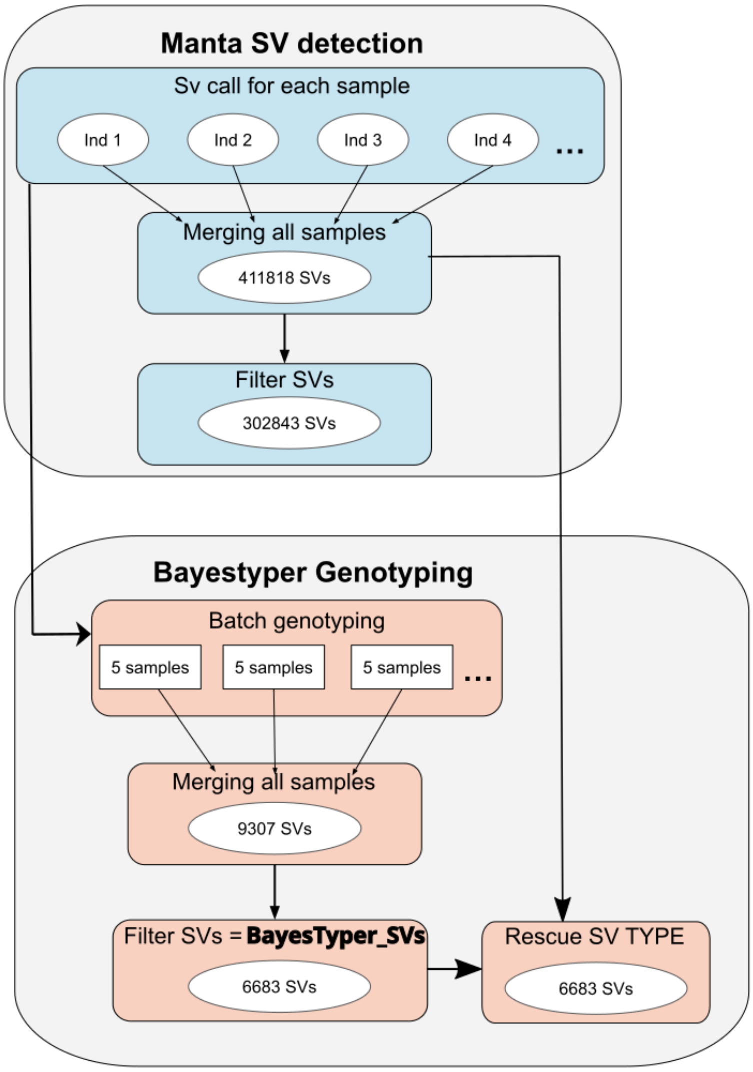


**Supplementary Figure 1: BayesTyper_SVs detection pipeline**

Top: Manta-based detection, bottom: BayesTyper genotyping. The final set BayesTyper_SVs used in this study is indicated in bold.

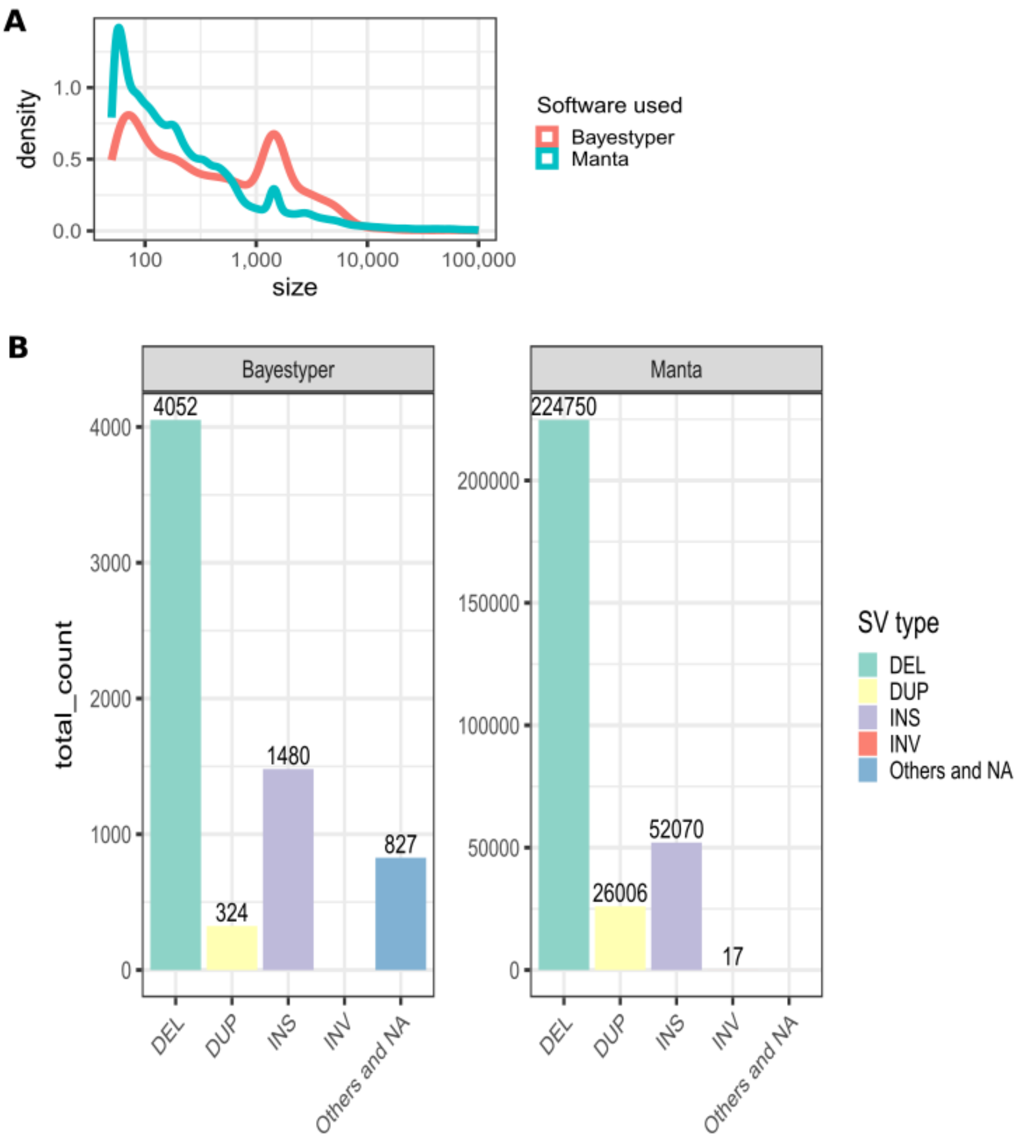


**Supplementary Figure 2: Comparison of SVs between manta and bayestyper**

**A:** Density of SV size (in log10). **B:** Number of SV of different types. SV types for bayestyper are not directly present but are rescued as described in **Supplementary Figure 1** and Material and methods.

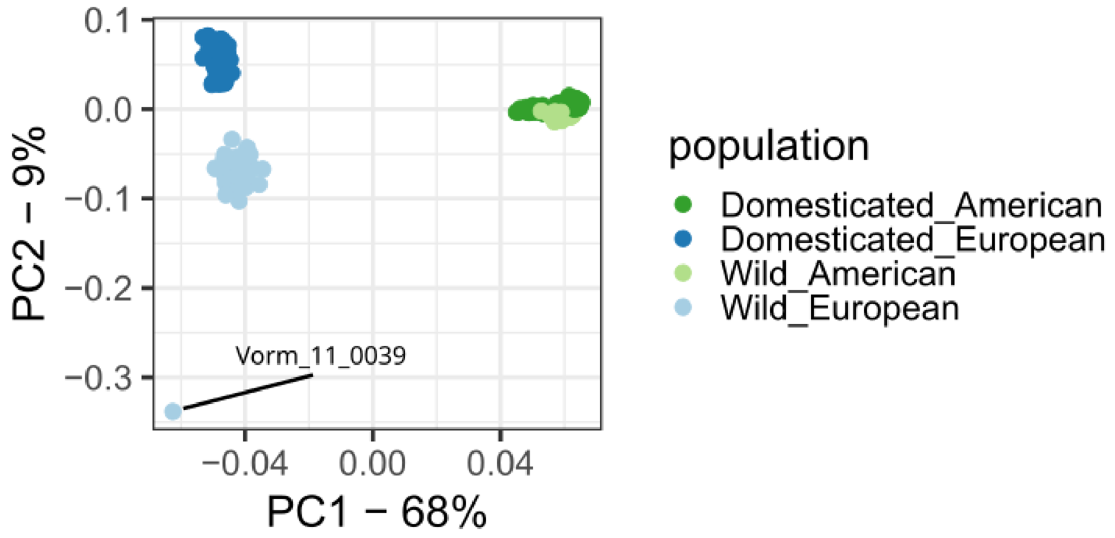


**Supplementary Figure 3: Whole genome PCA using BayesTyper_SVs**

PCA based on BayesTyper_SVs distinguish continents on PC1, and domestication status on PC2. The outlier individual (Vorm_11_0039 – Wild European) is indicated. While investigating this outlier by doing chromosome by chromosome PCA, we observed that the same individual was an outlier in 16 out of 29 chromosomes.

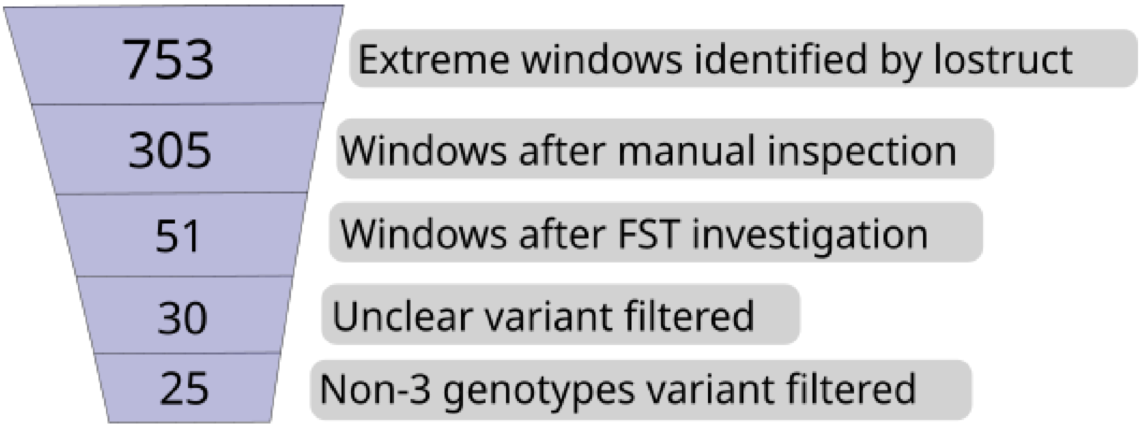


**Supplementary Figure 4: lostruct_SVs detection method pipeline**

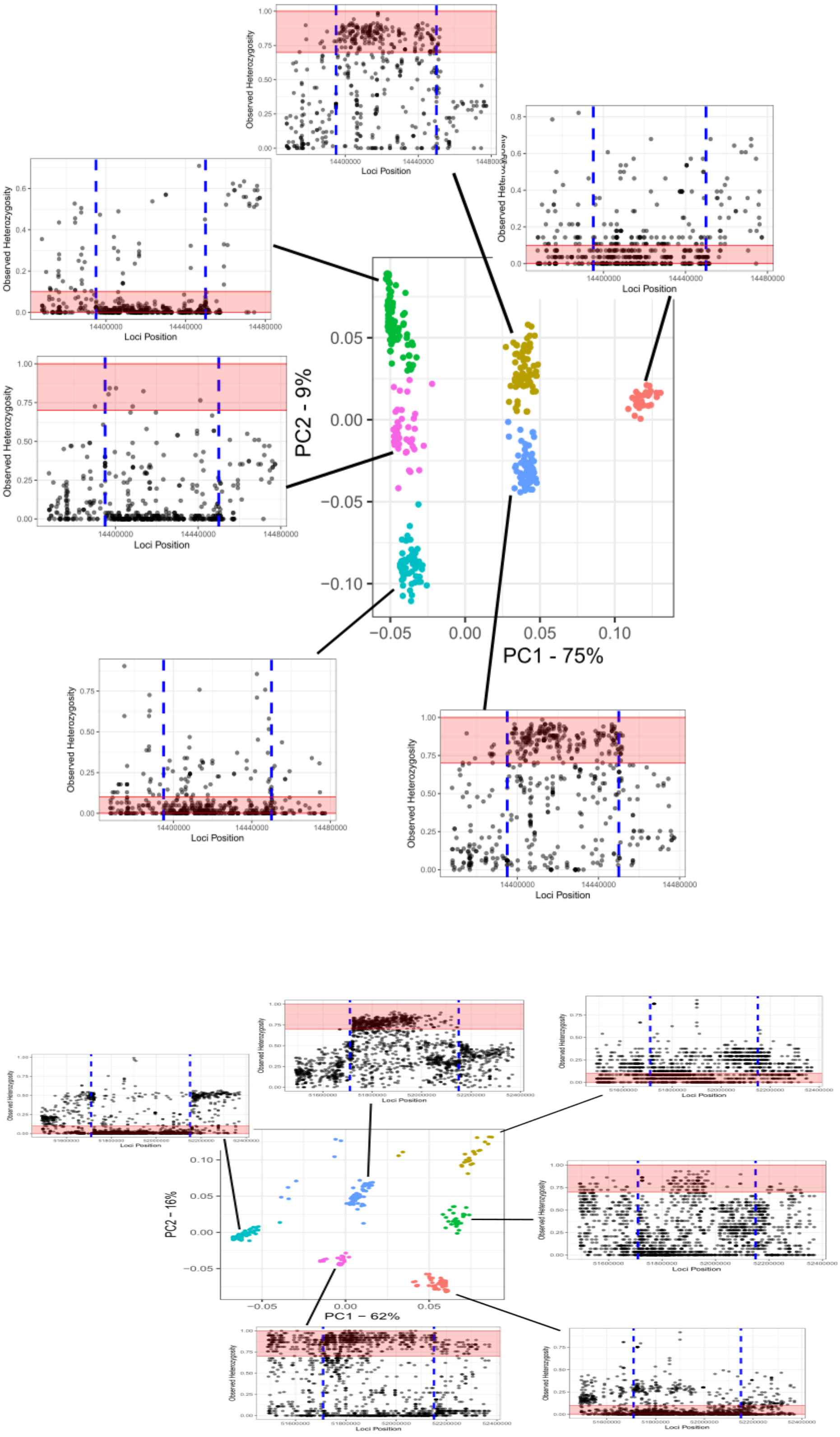


**Supplementary figure 5: Heterozygosity within PCA cluster**

Top : ssa02: 14395000-14450000 ; bottom: ssa13: 51710000-52150000. Red-shaded area indicates heterozygosity expectations in the case of an SV with 3 alleles (See **Figure 4D**).

Vertical dotted lines indicate the SV boundar

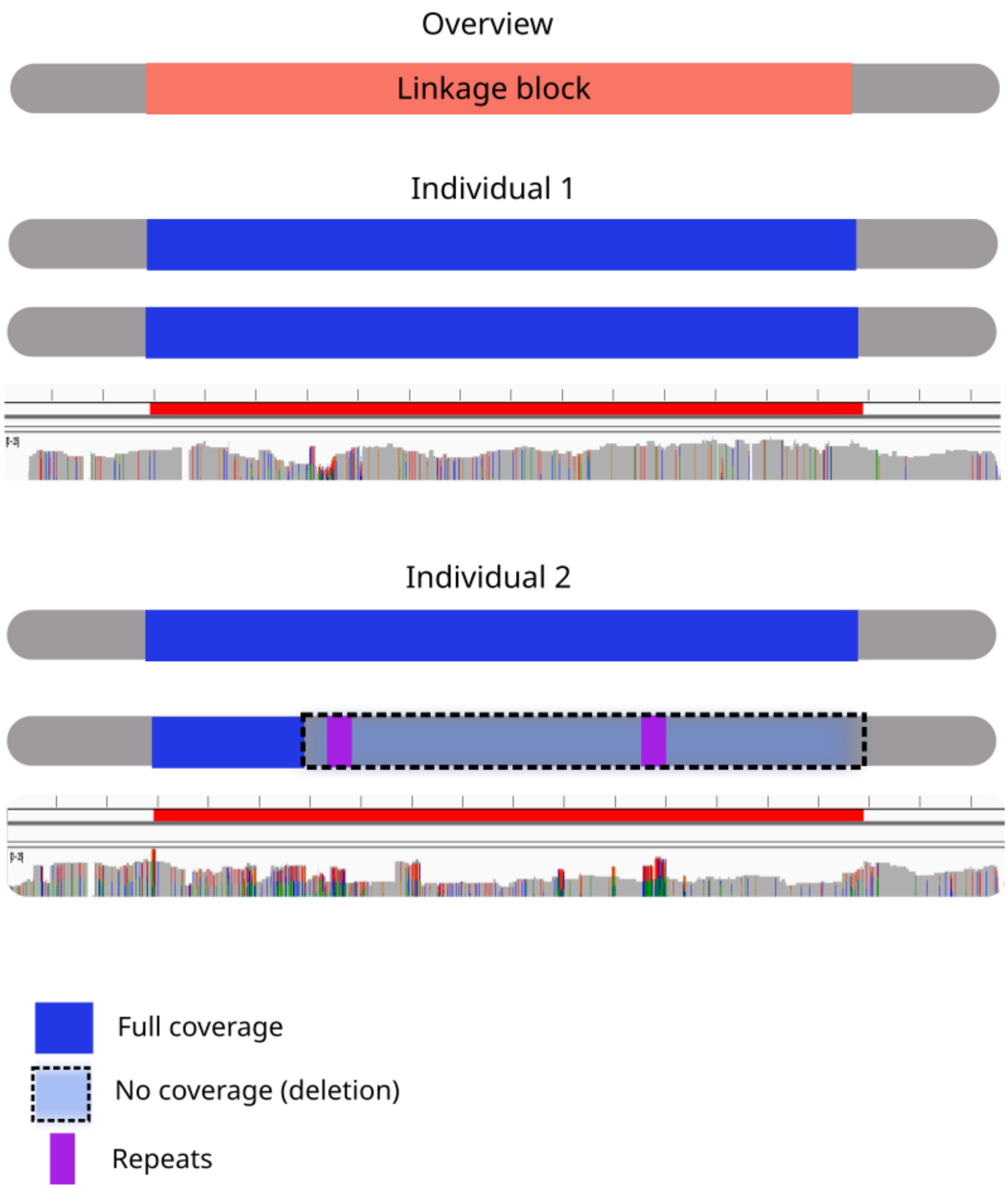


**Supplementary figure 6: IG_V_region coverage and putative deletion**

Two diploid individuals are represented, as well as an IGV screen of their sequencing depth (from 0 to 25) using long reads sequencing (see methods). On top, one individual has both chromosomes with full coverage. On bottom, the IGV screen indicates a half-drop (coherent with a heterozygote) in coverage for a part of the region, except in small regions with a very high genetic diversity, potentially indicating collapsed reads due to repeats.

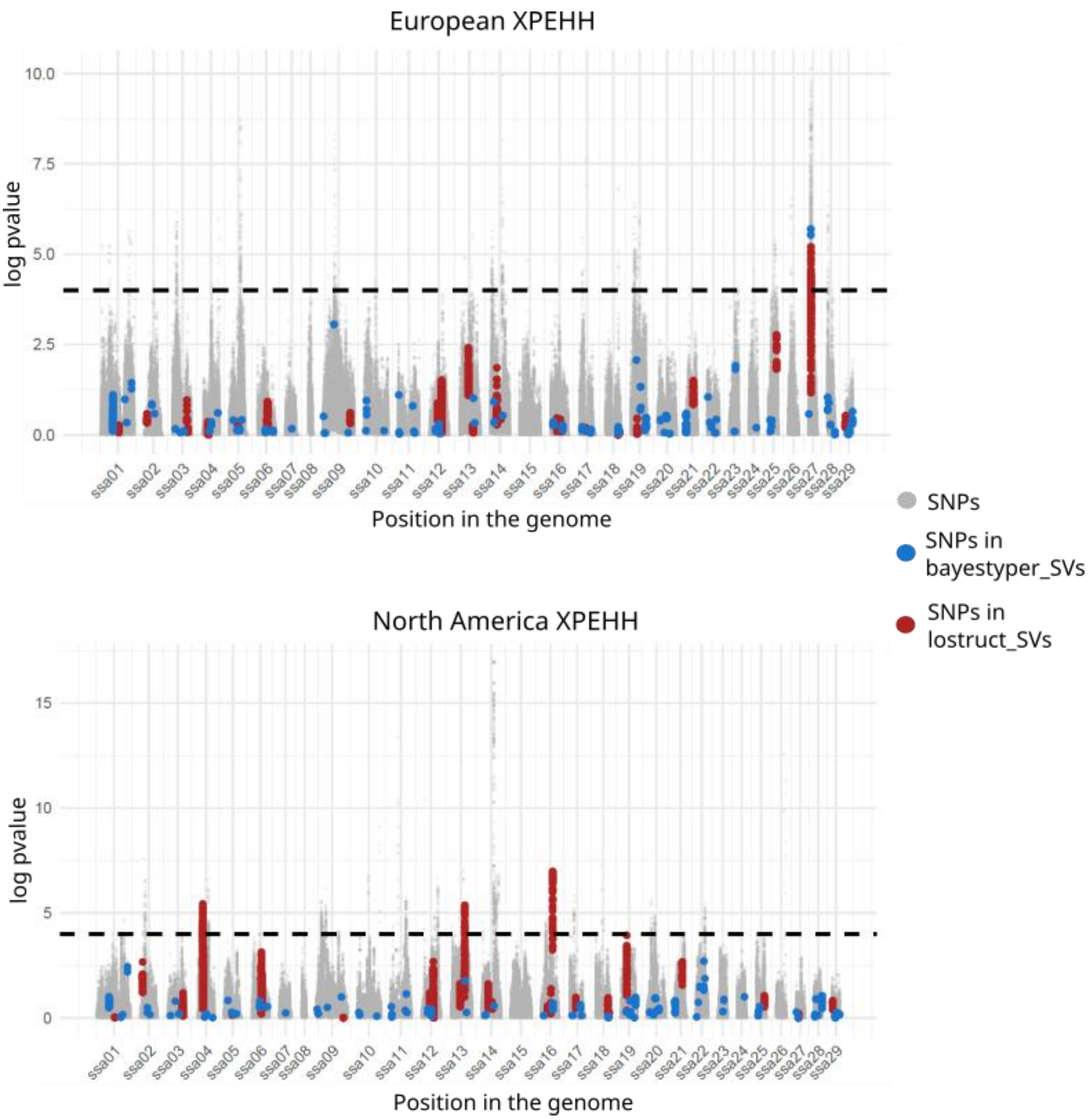


**Supplementary Figure 7: Genome-wide distribution of XP-EHH signals and overlap with structural variants**

Genome-wide distribution of XP-EHH scores across chromosomes, based on SNP data from Buso *et al.*, 2025. Grey points represent all SNPs included in the XP-EHH analysis. Colored points indicate SNPs overlapping with structural variants, with blue representing SNPs overlapping BayesTyper_SVs and red representing SNPs overlapping lostructs_SVs. The horizontal dotted line indicates the XP-EHH threshold value, as defined in Buso *et al.*, 2025, used to identify outlier signals.

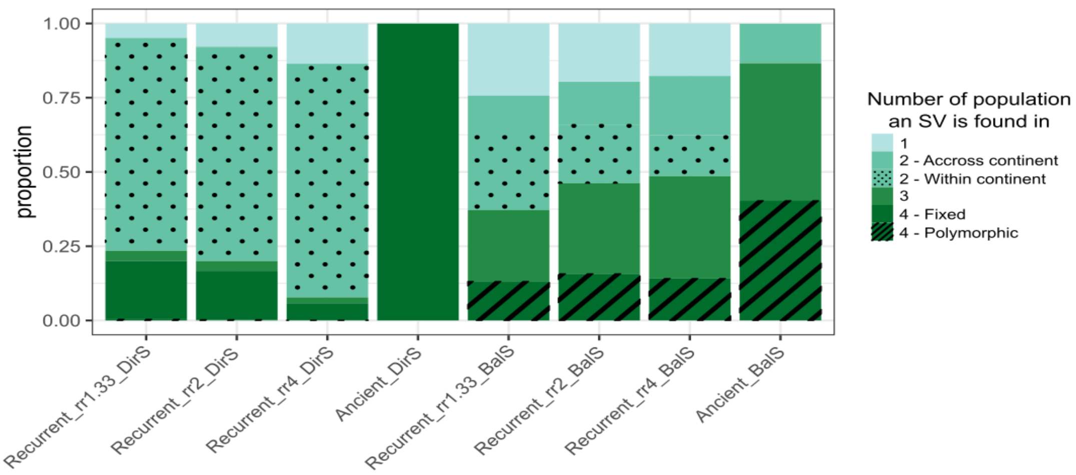


**Supplementary figure 8: Distribution of SVs across populations under different simulation scenarios.**

Stacked bar plots show the proportion of structural variants (SVs) observed in different populations, stratified by scenario. Labels indicate the evolutionary history of SVs (recurrent or ancient), the mode of selection (directional selection, DirS; balancing selection, BalS), and the recurrence rate (rr2, corresponding to an average of two SVs per simulation). All simulations were performed using a selection coefficient of 1.1.

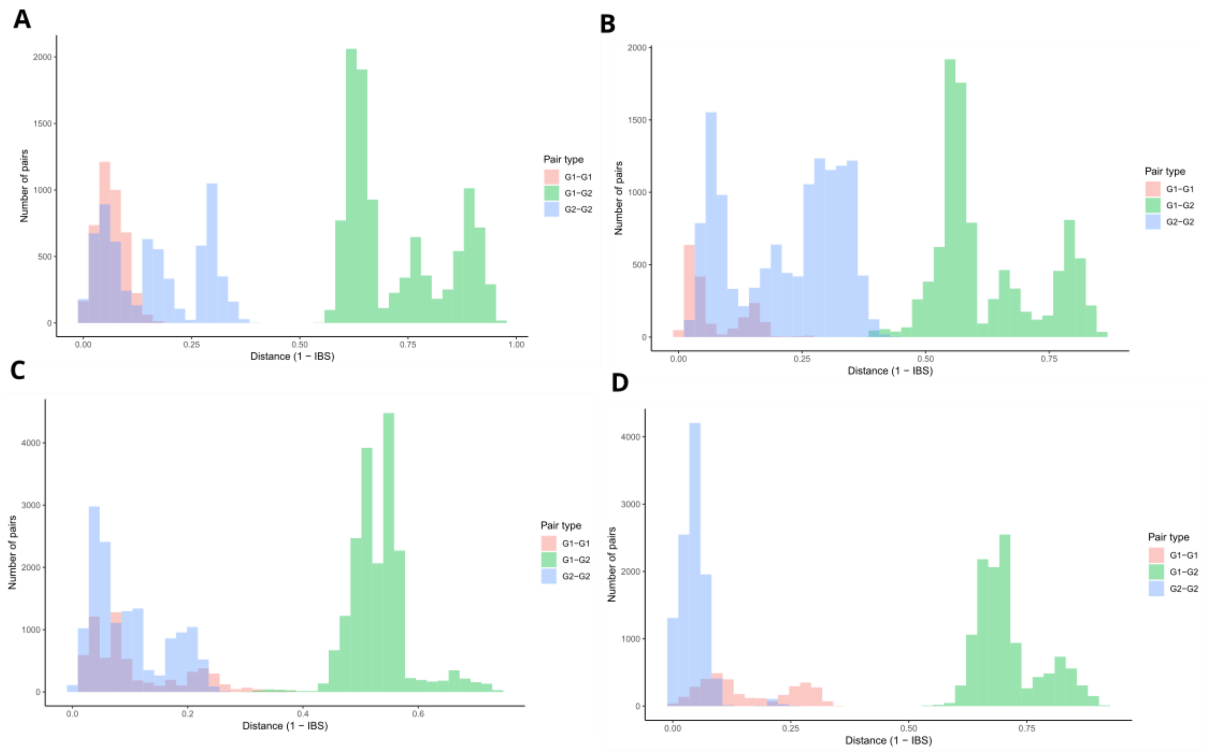


**Supplementary figure 9: Recurrent Lostruct_SVs.**
Genetic distance between SNPs in haplotype, G1-G1: AA vs AA; G1-G2:AA vs BB; G2-G2: BB vs BB. Under recurrent SV, the distribution is expected to be multimodal for G1-G2. A: ssa05:44020000-44084228; B: ssa16:33750000-34020915; C: ssa16:60620000-60880000; D: ssa21:38333640-38459102

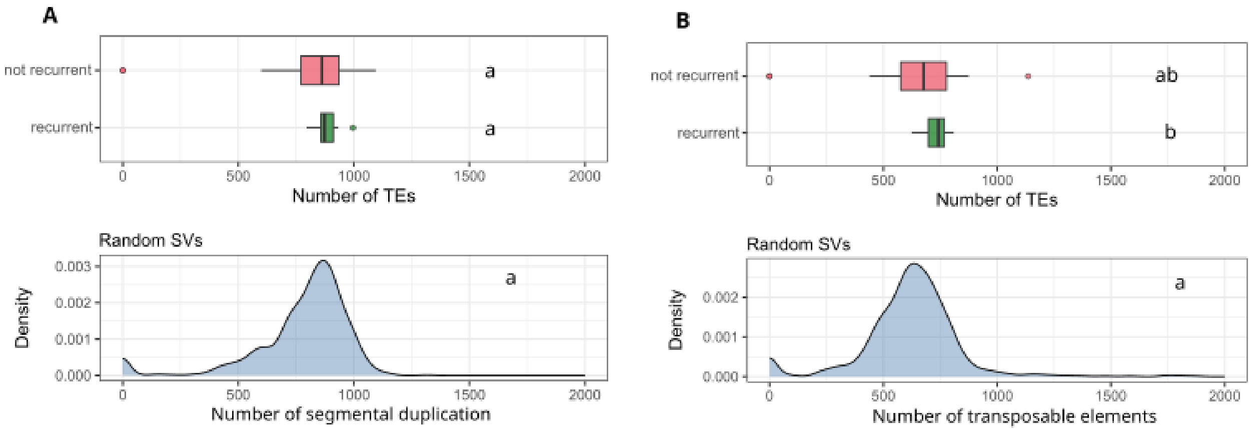


**Supplementary Figure 10 Transposable elements enrichment near Lostructs_SVs**

Number of repeats near SVs boundaries (+/- 100kb) for the 4 SVs assigned as recurrent (**Supplementary Figure 9**) or the other Lostruct_SVs. These numbers are compared to the numbers of repeat for 1000 randomly picked location in the genome (+/- 100kb). Letters indicate the significantly different groups **A:** Segmental duplications (not recurrent-recurrent: p-value=0.48; not recurrent-random: p-value=0.13; recurrent-random: p-value=0.15) **B:** Transposable elements (not recurrent-recurrent: p-value=0.13; not recurrent-random: p-value=0.10; recurrent-random: p-value=0.04)

**Supplementary table 1: Lostruct_SVs coordinates and overlapping genes**

| Coordinate | Genes |
|---|---|
| Ssa01: 92750000-92880000 | |
| Ssa02: 14395000-14450000 | ENSSSAG00000041032;ENSSSAG00000045818;ENSSSAG00000040824; ENSSSAG00000047726 |
| Ssa03: 75839027-75979106 | ENSSSAG00000001726;ENSSSAG00000001727;ENSSSAG00000099337; ENSSSAG00000095529;ENSSSAG00000083867;ENSSSAG00000090150; ENSSSAG00000004153;ENSSSAG00000057854;ENSSSAG00000009076; ENSSSAG00000004118 |
| Ssa03: 81220000-81261255 | ENSSSAG00000000943;ENSSSAG00000000925;ENSSSAG00000088970 |
| Ssa04: 29000000-32100000 | ENSSSAG00000005991;ENSSSAG00000041689;ENSSSAG00000109184; ENSSSAG00000113463;ENSSSAG00000045683;ENSSSAG00000086905; ENSSSAG00000049824;ENSSSAG00000049858;ENSSSAG00000089981; ENSSSAG00000111565;ENSSSAG00000059348;ENSSSAG00000041070; ENSSSAG00000005892;ENSSSAG00000120002;ENSSSAG00000005936; ENSSSAG00000005953;ENSSSAG00000041666;ENSSSAG00000041774; ENSSSAG00000103400;ENSSSAG00000043249;ENSSSAG00000111674; ENSSSAG00000043456;ENSSSAG00000045334;ENSSSAG00000045653; ENSSSAG00000010182;ENSSSAG00000005502;ENSSSAG00000045602; ENSSSAG00000084245;ENSSSAG00000047918;ENSSSAG00000107634; ENSSSAG00000049840;ENSSSAG00000085065;ENSSSAG00000036829; ENSSSAG00000040099;ENSSSAG00000041175;ENSSSAG00000041197; ENSSSAG00000005978;ENSSSAG00000006503;ENSSSAG00000043307; ENSSSAG00000063298;ENSSSAG00000096375;ENSSSAG00000045755; ENSSSAG00000049796;ENSSSAG00000088225;ENSSSAG00000102949; ENSSSAG00000049978;ENSSSAG00000054868;ENSSSAG00000054962; ENSSSAG00000057700;ENSSSAG00000041051;ENSSSAG00000041721; ENSSSAG00000049715;ENSSSAG00000049619 |
| Ssa05:44020000-44084228 | |
| Ssa06: 52540000-55000000 | ENSSSAG00000100736;ENSSSAG00000032979;ENSSSAG00000107853; ENSSSAG00000003730;ENSSSAG00000003760;ENSSSAG00000007969; ENSSSAG00000104602;ENSSSAG00000094877;ENSSSAG00000067479; ENSSSAG00000120792;ENSSSAG00000120858;ENSSSAG00000007629; ENSSSAG00000114918;ENSSSAG00000007615;ENSSSAG00000002811; |

| | |
|---|---|
| | ENSSSAG00000115651;ENSSSAG00000003941;ENSSSAG00000100701; ENSSSAG00000008124;ENSSSAG00000015636;ENSSSAG00000087136; ENSSSAG00000067588;ENSSSAG00000072139;ENSSSAG00000121038; ENSSSAG00000007650;ENSSSAG00000007605;ENSSSAG00000103681; ENSSSAG00000022058;ENSSSAG00000072133;ENSSSAG00000088042; ENSSSAG00000094083 |
| Ssa09: 140185732-140202636 | ENSSSAG00000057167;ENSSSAG00000057206 |
| Ssa12: 63150000-63800000 | ENSSSAG00000069806;ENSSSAG00000077210;ENSSSAG00000119026; ENSSSAG00000069809;ENSSSAG00000094905;ENSSSAG00000077213; ENSSSAG00000079782;ENSSSAG00000097016;ENSSSAG00000070160; ENSSSAG00000098839 |
| Ssa12: 64000000-65000000 | ENSSSAG00000082416;ENSSSAG00000084198;ENSSSAG00000094754; ENSSSAG00000080980;ENSSSAG00000080100;ENSSSAG00000080864; ENSSSAG00000104465;ENSSSAG00000080857;ENSSSAG00000080876; ENSSSAG00000080924;ENSSSAG00000080995;ENSSSAG00000081022; ENSSSAG00000081065;ENSSSAG00000084657;ENSSSAG00000080247; ENSSSAG00000080851;ENSSSAG00000008488;ENSSSAG00000081034; ENSSSAG00000081196 |
| Ssa13: 51710000-52150000 | ENSSSAG00000111810;ENSSSAG00000070051;ENSSSAG00000070458; ENSSSAG00000073513 |
| Ssa13: 76000000-76800000 | ENSSSAG00000109016;ENSSSAG00000058260;ENSSSAG00000100170; ENSSSAG00000058310;ENSSSAG00000102970;ENSSSAG00000058323; ENSSSAG00000058336;ENSSSAG00000058211 |
| Ssa14: 35000000-35400000 | ENSSSAG00000022880;ENSSSAG00000090634;ENSSSAG00000117177; ENSSSAG00000032035;ENSSSAG00000048190;ENSSSAG00000101138; ENSSSAG00000048567;ENSSSAG00000028979;ENSSSAG00000048728 |
| Ssa14: 56242199-56291923 | ENSSSAG00000055927 |
| Ssa16: 33750000-34020915 | ENSSSAG00000050779;ENSSSAG00000069422;ENSSSAG00000108830; ENSSSAG00000069428 |
| Ssa16: 51650000-51820000 | ENSSSAG00000007201;ENSSSAG00000007205;ENSSSAG00000007185; ENSSSAG00000121435;ENSSSAG00000112886;ENSSSAG00000006625 |
| Ssa16: 60620000-60880000 | ENSSSAG00000086157;ENSSSAG00000039914 |
| Ssa17: 35400419-35510000 | ENSSSAG00000111008;ENSSSAG00000110934;ENSSSAG00000111335; ENSSSAG00000083564;ENSSSAG00000111627 |
| Ssa18: 67200000-67510731 | |
| Ssa18: 72307638-72471162 | ENSSSAG00000066453;ENSSSAG00000066459;ENSSSAG00000101816; ENSSSAG00000040226;ENSSSAG00000102714;ENSSSAG00000040378; ENSSSAG00000040458;ENSSSAG00000040529;ENSSSAG00000092702; ENSSSAG00000040184;ENSSSAG00000040291;ENSSSAG00000040351; ENSSSAG00000062190;ENSSSAG00000112668;ENSSSAG00000062249; ENSSSAG00000103344;ENSSSAG00000062102;ENSSSAG00000117465 |
| Ssa19: 31600000-32200000 | ENSSSAG00000072920;ENSSSAG00000072912;ENSSSAG00000072943; ENSSSAG00000115609;ENSSSAG00000091716 |
| Ssa21: 38333640-38459102 | ENSSSAG00000059086;ENSSSAG00000059799 |
| Ssa25: 43856943-43882324 | ENSSSAG00000114775 |
| Ssa27: 11940000-12240000 | ENSSSAG00000068404;ENSSSAG00000082142;ENSSSAG00000040147; ENSSSAG00000040125;ENSSSAG00000039984;ENSSSAG00000039965; ENSSSAG00000084331;ENSSSAG00000120981;ENSSSAG00000107011; ENSSSAG00000115878;ENSSSAG00000068332 |
| Ssa29: 5670000-5900000 | |

**Supplementary table 2: Genes in the IG_V_region**

Genes overlapping the IG_V_region, based on Ssal_v3.1 annotations. NCBI annotations are retrieved from near-overlapping position of NCBI and ensembl annotation done in a genome browser tool. Other annotations are indicated with the methods used (either blast against zebrafish or from gene sequence proximity with closely related species using the gene tree tools in ensembl)

| ENSEMBL gene code | ENSEMBL annotation or biotype | NCBI gene code | NCBI annotation or biotype | Others annotations (source) |
|---|---|---|---|---|
| ENSSSAG00000004118 | Protein coding | LOC123741723 | lncRNA | - |

| | | | | |
|---|---|---|---|---|
| ENSSSAG00000001726 | IG_V_gene | - | - | Immunoglobulin (blast against zebrafish) |
| ENSSSAG00000001727 | IG_V_gene | LOC106601724 | V_segment | Immunoglobulin (blast against zebrafish) |
| ENSSSAG00000099337 | IG_V_gene | LOC123741721 | lncRNA | Immunoglobulin (blast against zebrafish) |
| ENSSSAG00000095529 | Protein coding | - | - | Immunoglobulin (blast against zebrafish) |
| ENSSSAG00000009076 | Protein coding | - | - | Immunoglobulin (Closely related sequence to pike) |
| ENSSSAG00000083867 | IG_V_gene | - | - | Immunoglobulin (blast against zebrafish) |
| ENSSSAG00000090150 | Protein coding | - | - | Rho GTPase activating protein 25 *arhgap25* (blast against zebrafish) |
| ENSSSAG00000004153 | IG_V_gene | LOC106601682 | V_segment | Immunoglobulin (blast against zebrafish) |
| ENSSSAG00000057854 | IG_V_gene | LOC106592589 | V_segment | Immunoglobulin (closely related sequence to brown trout) |

**Supplementary table 3: Genes overlapping structural variants highlighted in Figure S6**

List of annotated genes located within structural variant (SV) regions that overlap with XP-EHH outlier signals shown in **Supplementary figure 7**. Gene identifiers, gene names, and corresponding SV genomic coordinates are shown.

| Gene | Gene name | SV coordinate |
|---|---|---|
| ENSSSAG00000038615 | TatD DNase domain containing 1 (*tatdn1*) | ssa27: 11940000-12240000 |
| ENSSSAG00000068332 | GA binding protein transcription factor subunit beta 2a (*gabpb2a*) | ssa27: 11940000-12240000 |
| ENSSSAG00000115878 | Unknown | ssa27: 11940000-12240000 |
| ENSSSAG00000107011 | guanine nucleotide-binding protein G(s) subunit alpha-like | ssa27: 11940000-12240000 |
| ENSSSAG00000120981 | Unknown | ssa27: 11940000-12240000 |
| ENSSSAG00000084331 | MLLT11 transcription factor 7 cofactor | ssa27: 11940000-12240000 |
| ENSSSAG00000039965 | CDC42 small effector 1 (*CDC42SE1*) | ssa27: 11940000-12240000 |
| ENSSSAG00000039984 | BCL2/adenovirus E1B 19 kDa protein-interacting protein 2 | ssa27: 11940000-12240000 |
| ENSSSAG00000040125 | Prune exopolyphosphatase 1 (*PRUNE1*) | ssa27: 11940000-12240000 |
| ENSSSAG00000040147 | MINDY lysine 48 deubiquitinase 1 (*MINDY1*) | ssa27: 11940000-12240000 |
| ENSSSAG00000082142 | DNA-binding protein RFX5-like | ssa27: 11940000-12240000 |
| ENSSSAG00000068404 | phosphatidylinositol 4-kinase beta | ssa27: 11940000-12240000 |
| ENSSSAG00000005991 | Helt bHLH transcription factor (*helt*) | ssa04:29000000_32100000 |
| ENSSSAG00000005936 | Cilia and flagella associated protein 97 (*cfap97*) | ssa04:29000000_32100000 |

| | | |
|---|---|---|
| ENSSSAG00000005953 | Solute carrier family 25 member 4 (*slc25a4*) | ssa04:29000000_32100000 |
| ENSSSAG00000005978 | EF-hand calcium-binding domain-containing protein 6 | ssa04:29000000_32100000 |
| ENSSSAG00000006503 | Acyl-CoA synthetase long chain family member 1a (*acsl1a*) | ssa04:29000000_32100000 |
| ENSSSAG00000058310 | CCR4-NOT transcription complex subunit 8 (*cnot8*) | ssa13:76000000_76800000 |
| ENSSSAG00000102970 | Gem (nuclear organelle) associated protein 5 (*gemin5*) | ssa13:76000000_76800000 |
| ENSSSAG00000058323 | Family with sequence similarity 114 member A2 (*FAM114A2*) | ssa13:76000000_76800000 |
| ENSSSAG00000039914 | Cholecystokinin B receptor a (*cckbra*) | ssa16: 60620000_60880000 |
